# The role of vegetative cell fusions in the lifestyle of the wheat fungal pathogen *Zymoseptoria tritici*

**DOI:** 10.1101/2020.01.26.918797

**Authors:** Carolina Sardinha Francisco, Maria Manuela Zwyssig, Javier Palma-Guerrero

## Abstract

The ability of fungal cells to undergo cell fusion allows them to maximize their overall fitness. In this study, we characterized the role of the *so* gene orthologous in *Zymoseptoria tritici* and the biological contribution of vegetative cell fusions in the lifestyle of this latent necrotrophic fungus. Firstly, we show that *Z. tritici* undergoes self-fusion between distinct cellular structures and its mechanism is dependent on the initial cell density. Next, the deletion of *ZtSo* resulted in the loss of cell-to-cell communication affecting both hyphal and germlings fusion. We show that *Z. tritici* mutants for MAP kinase-encoding *ZtSlt2* (orthologous MAK-1) and *ZtFus3* (orthologous MAK-2) genes also fail to undergo self-stimulation and self-fusion, demonstrating the functional conservation of this signaling mechanism across species. Additionally, the *ΔZtSo* mutant was severely impaired in melanization, which leads us to identify a trade-off between fungal growth and melanization. Though it has been proposed that So is a scaffold protein for MAP kinase genes from the CWI pathway, its deletion did not affect the cell wall integrity of the fungus. Finally, we demonstrated that anastomose is dispensable for pathogenicity, but essential for the fruiting body development and its absence abolish the asexual reproduction of *Z. tritici*. Taken together, our data show that *ZtSo* is required for fungal development, while vegetative cell fusions are essential for fungal fitness.

## Introduction

Communication is a ubiquitous primitive characteristic developed by all living species. From high mammals to the simplest forms of life, the exchange of signals between organisms have driven the evolution of the species (Endler, 1993, Ord & Garcia-Porta, 2012, Wilson, 1975). The ability to communicate effectively may affect mating, predation, competition, dominance hierarchy, signal modalities, and survival (Endler, 1993, Gillam, 2011, Wilson, 1975). This complex mechanism starts when a given organism (the sender) secretes in the environment a self-produced molecular signal (the message) which alters the behavior of another organism (the receiver) (Endler, 1993, Wilson, 1975). Communication also happens at the cellular level. This so-called “cell-to-cell communication” creates a complex signaling network that involves different extracellular signals and distinct cell types that regulate manifold pathways (van Gestel *et al.*, 2012, Shrout *et al.*, 2011, Fischer & Glass, 2019). Inter- and intraspecies cell-to-cell communication has been widely studied in fungi to address biological functions including the secretion of pheromones to attract the opposite sexual partner; the production of quorum sensing molecules controlling the expression of virulence factors and morphological changes and, the regulation of cell fusions during vegetative growth (Cottier & Muhlschlegel, 2012, Bloemendal & Kuck, 2013, Khang *et al.*, 2010, Wongsuk *et al.*, 2016, Fischer & Glass, 2019).

The fungal mycelium is formed by three integrated processes including hyphal extension, branching and vegetative hyphal fusion -VHF (also known as anastomosis) (Glass *et al.*, 2000). In this last-mentioned process, two growing cells with identical vegetative compatibility loci engage in cell-to-cell communication, known as “ping-pong self-signaling”, which is thought to involve the secretion of unknown diffusible molecules and results in re-direct polarized hyphal growth toward each other. After physical contact, the cell walls are remodeled, the plasma membranes fused and the two interconnected cells exchange cytoplasm and organelles (Fleissner *et al.*, 2009). Whether the anastomosed individuals are vegetatively incompatible, the two fused cells rapidly collapse following DNA degradation by programmed cell death or they are severely inhibited in their growth (Saupe, 2000). Albeit non-self-anastomoses are described (Roca *et al.*, 2004, He *et al.*, 1998), this might be a very rare event in nature. It is widely accepted that mycelial network formed through VHF facilitates the intra-hyphal communication, translocation of water, nutrients, and signal molecules, which improves the general homeostasis and the spatial expansion of the fungal colony (Read *et al.*, 2010, Hickey *et al.*, 2002). In some pathogenic fungi, hyphal fusion is required for pathogenicity and host adhesion (Craven *et al.*, 2008, Prados Rosales & Di Pietro, 2008). Fungal cell fusions can also occur between conidial cells. Conidium is the asexual spore of many Ascomycetes and Basidiomycetes species. The process of fusion between germinating conidia involves the formation and interaction of specialized hyphae, called conidial anastomosis tube (CAT). CAT is thinner and shorter than VHF, and its induction dependent on nutrient deprivation and initial cell density (Roca *et al.*, 2005a). It has been postulated that CATs improve colony establishment, as well as they may increase the genetic variability by facilitating heterokaryosis and parasexual recombination, especially for those fungal species lacking sexual reproduction (Roca et al., 2005a). Evidence of gene and chromosome transfer between intra- or inter-fungal species has been shown as a mechanism to acquire pathogenicity or to broaden the host specificity (Mehrabi *et al.*, 2011, Friesen *et al.*, 2006, Temporini & VanEtten, 2004). On the other hand, anastomose can also be high-risk gambling by the acquisition of infectious cytoplasmic and genetic elements, such as mycoviruses, selfish elements or debilitated organelles, that can multiply uncontrolled and drain the energetic sources of the cell (Saupe, 2000, Biella *et al.*, 2002).

In the last decades, different studies about the molecular mechanisms underlying cell fusion identified several mutants defective in anastomosis, revealing that fungal communication and fusion are complex mechanisms, which encompasses components of several signaling pathways (Fu *et al.*, 2011, Xiang *et al.*, 2002, Aldabbous *et al.*, 2010, Pandey *et al.*, 2004, Fu *et al.*, 2014, Fischer & Glass, 2019). Hitherto, the best-characterized mutant is for the *soft* (*so*) gene of *Neurospora crassa*, a gene from the mitogen-activated protein (MAP) kinase MAK-1 pathway, which encodes a protein with unknown function (Fleissner *et al.*, 2005). The So protein is proposed to be the scaffold for cell wall sensors belonging to the MAP kinase cascade from the cell wall integrity (CWI) signaling pathway (Fischer & Glass, 2019). Furthermore, the *so* gene has been shown to have an essential role in the hyphal anastomosis, presumably by regulating the secretion of an undefined chemoattractant in an oscillatory manner with the MAK-2 from the MAP kinase signal response pathway (Fleissner et al., 2009). Beyond *N. crassa*, the *so* gene has been characterized in different filamentous fungi including model organisms, plant pathogens, and endophytic fungi (Engh *et al.*, 2007, Maruyama *et al.*, 2010, Craven et al., 2008, Prados Rosales & Di Pietro, 2008, Charlton *et al.*, 2012). Though all *so* mutants fail to undergo hyphal fusions, the distinct effects on pathogenicity reported by the deletion of the *so* gene in different fungi suggests that the biological contribution of anastomosis might depend on the infection strategies developed by different fungal pathogen species.

*Z. tritici* is an apoplastic pathogen with a latent necrotrophic lifestyle and considered the most damaging pathogen of wheat in Europe (Fones & Gurr, 2015). Hyphae formed from either germinated ascospores (sexual spore), pycnidiospores (asexual spores) or blastospores (asexual spores produced by budding) are essential for penetrating wheat leaves through stomata and colonization of the apoplastic space. After a long asymptomatic phase (which varies depending on the wheat genotype and fungal strain combination), the onset of the necrotrophic phase is followed by the appearance of lesions, disintegration of host tissue and formation of asexual fruiting bodies. Though *Z. tritici* is among the top 10 most studied phytopathogens (Dean *et al.*, 2012), little is known about vegetative cell fusion in this organism. To date, it has been shown that the deletion of the β-subunit of the heterotrimeric G protein - *MgGpb1* or the *ZtWor1*, a transcriptional regulator of genes located downstream of the cyclic adenosine monophosphate (cAMP) pathway, negatively regulate cell fusion producing germ tubes that undergo extensive anastomosis (Mehrabi *et al.*, 2009, Gohari *et al.*, 2014).

In this study, we aimed to determine whether the *ZtSo* gene and vegetative cell fusions play important biological roles in the lifestyle of *Z. tritici*. We showed that the ubiquitous ability of *Z. tritici* to undergo self-fusion was disrupted by the deletion of *ZtSo* affecting both hyphal and germling fusions. The characterization of mutants lacking the MAP kinase-encoding genes *ZtSlt2* (orthologous MAK-1) or *ZtFus3* (orthologous MAK-2) showed that the cell fusion of *Z. tritici* is also regulated by the ping-pong self-signaling mechanism, demonstrating the functional conservation of this mechanism across species. We found that *ZtSo* is required for vegetative growth and melanization, but not to maintain the cellular integrity of the fungus. We discovered that anastomoses are dispensable for pathogenicity, but they are essential for fruiting body development. In the absence of cell fusions, *Z. tritici* does not undergo asexual reproduction. These findings illustrate the impact of *ZtSo* for fungal development and the importance of vegetative cell fusions for fungal fitness.

## Results

### Cell fusion in *Z. tritici* allows the bidirectional transfer of cytoplasmic content, but not nuclei exchange

We co-inoculated either blastospores or pycnidiospores of both 1E4_GFP_ and 1E4_mCh_ fluorescent strains onto water agar (WA - 1% agar in water), a hyphal-inducing medium, to investigate the ability of *Z. tritici* to undergo self-fusions. Fungal cells expressing each fluorescent protein appeared in only one color-channel of the fluorescence microscope (Fig. 1A), which ensures that the reciprocal cytoplasmic streaming is explicitly due to cell fusions. Though *Z. tritici* produces blastospore and pycnidiospores as asexual spores instead of conidium, we used the CAT terminology to define the fusion between germinating spores. CATs formed between blastospore or pycnidiospores germlings started after 4 hours of incubation, but are frequently observed after 17 hours of incubation (Fig. 1B). Vegetative hyphal fusions (VHFs) from germinated blastospores or pycnidiospores were noticed at 40 hours after incubation (hai) (Fig. 1C and D). Multiple interconnections via fusion bridges were observed in all tested morphotypes (Fig. 1B-D). The co-infection of wheat plants using either blastospores or pycnidiospores of 1E4_GFP_ and 1E4_mCh_ strains also resulted in VHFs on the wheat leaf surface (Fig. S1). Self-fusions and cytoplasmic mixing occurred in the first 48 hai (Fig. S1A-B). It indicates that VHFs may play a role during the initial fungal establishment and colonization of the host tissues.

**Figure 1.**
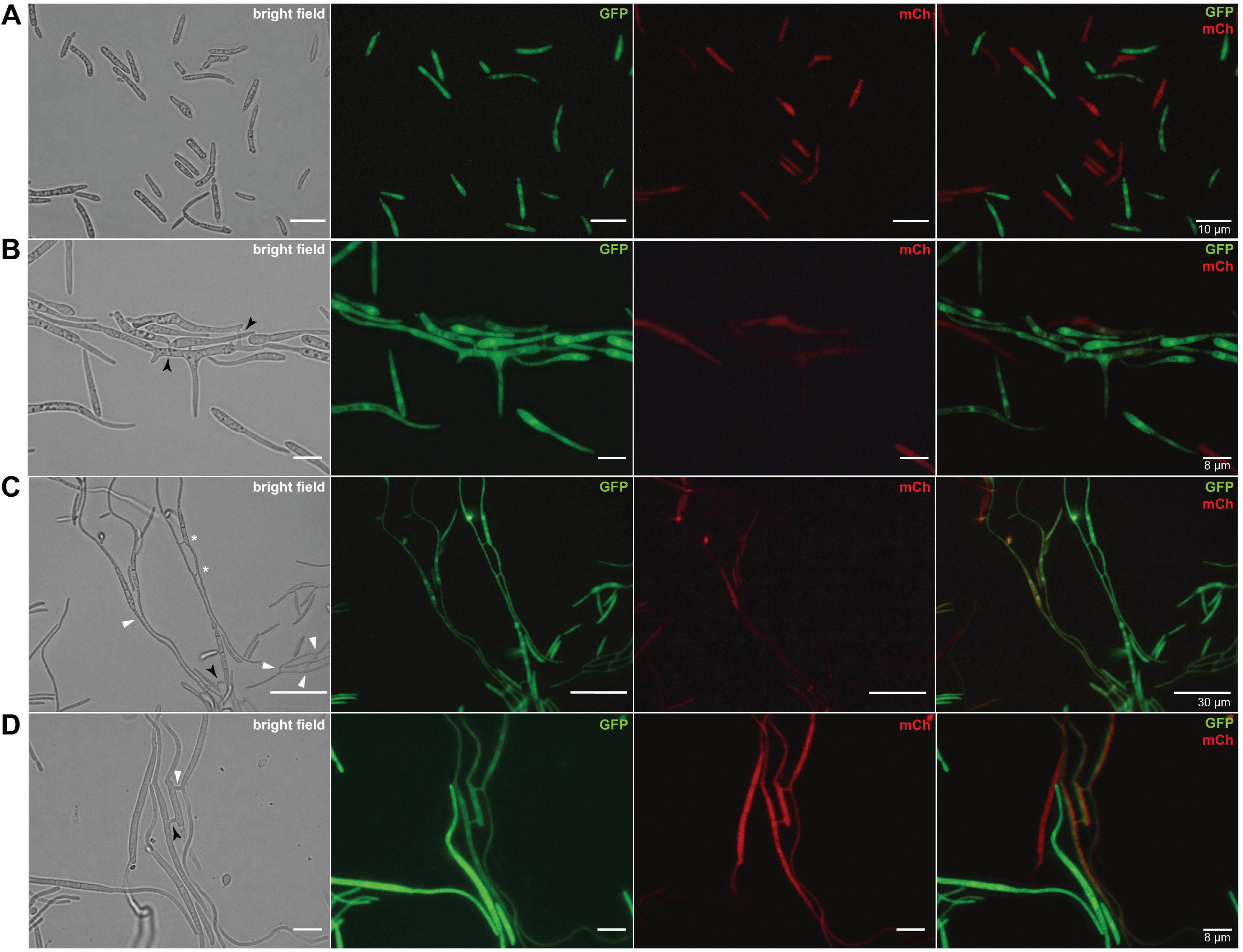
Vegetative cell fusions of *Zymoseptoria tritici*. Blastospores or pycnidiospores expressing either the cytoplasmic green fluorescent protein (GFP) or red-fluorescent protein (mCherry or mCh) were co-inoculated on water agar (WA) plates in a 1:1 ratio. (A) Blastospores-tagged GFP or mCh appeared in only one-color channel of the fluorescence microscope. (B) High initial cell concentration (1×10^7^ blastospore/mL) induces conidial anastomosis tubes (CATs) that were frequently observed after 17 hours of incubation. (C-D) Vegetative hyphal fusions (VHFs) from germinating blastospores or pycnidiospores are induced at lower initial cell concentration (1×10^6^ blastospore/mL) and were noticed after 40 hours of incubation, respectively. Black arrows or white triangles point to the CATs or VHFs, respectively, which resulted in the mixture of both GFP and mCh fluorescent proteins due to the exchange of cytoplasm content between the two fused cells. White asterisks point to the fusion bridges formed from individuals expressing the same fluorescent proteins.

To monitor the genetic exchange between genetically identical *Z. tritici*, we used the IPO323 ZtHis1-ZtGFP strain. We looked for the presence of multinucleated cells surrounding hyphal fusion points. None cell compartments containing more than one nucleus, nor a nuclei exchange between the two interconnected hyphae were observed (Fig. 2). After the fusion bridge formation, one of the neighbor’s nuclei divides and migrate to occupy the new septal compartment. This finding suggests that cell fusion in *Z. tritici* results only on cytoplasmic mixing, but not in heterokaryon formation or parasexuality.

**Figure 2.**
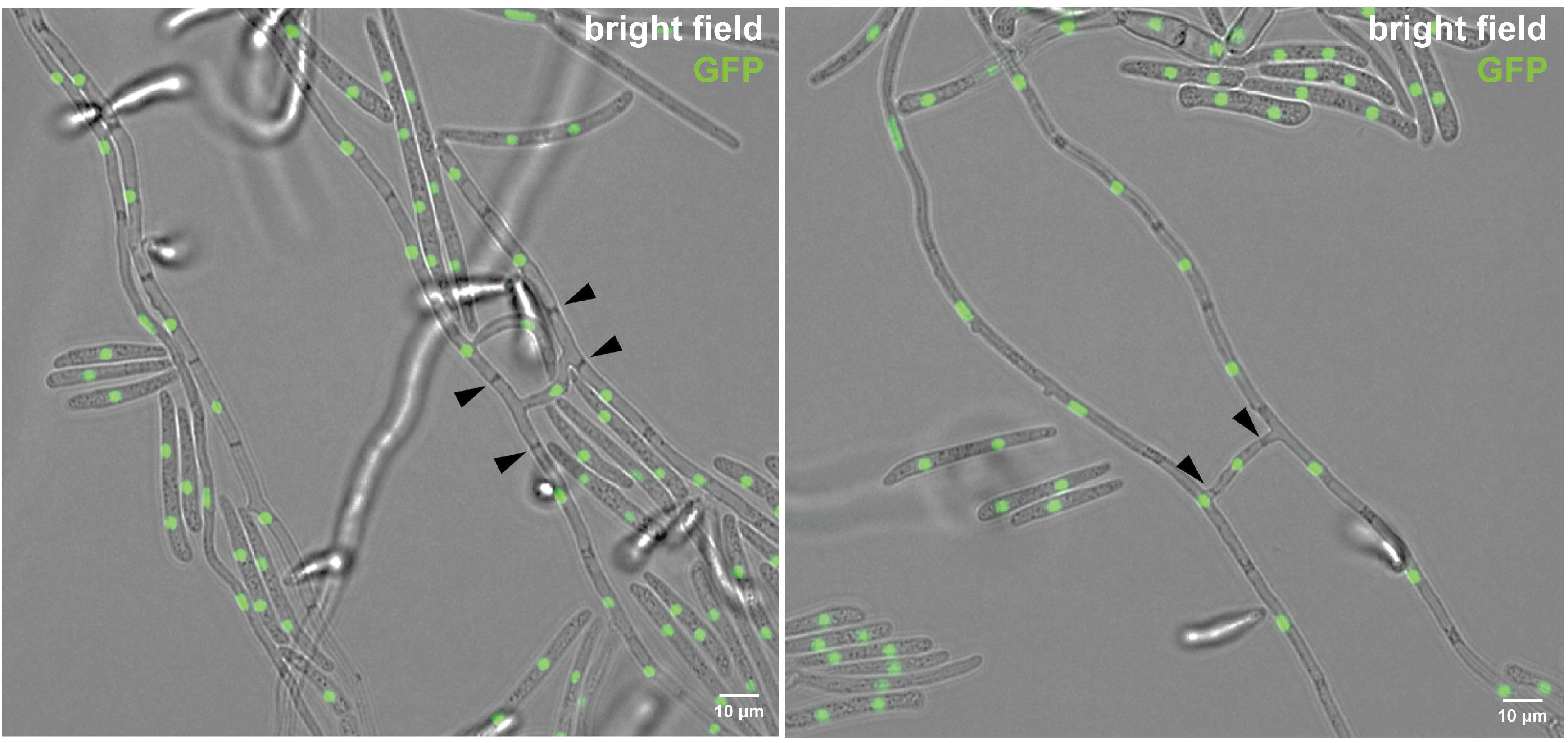
Hyphal fusion does not lead to the generation of heterokaryon individuals in *Zymoseptoria tritici*. ZtHis1-ZtGFP strain was used to monitor the nuclei movement during hyphal fusion. None septum containing more than one nucleus was observed, neither the genetic exchange between the two fused hyphal individuals. Black triangles indicate the septal compartment formed in the fusion bridges containing only one nucleus.

### Orthologues of *ZtSo* are widely distributed within the Dothideomycetes

We identified the *so* orthologous (*ZtSo*) in the *Z. tritici* genome. The *ZtSo* gene (*Mycgr3G74194* or *Zt09_7_00503*) consist of 3,794 bp open reading frame and encodes a polypeptide of 1,227 amino acid (Fig. S2A). Overall, proteins with varying degrees of similarity to ZtSo were identified among several Dothideomycetes species (n=20). We also found homology with other four fungal Classes, including Sordariomycetes (n=4), Xylomycetes (n=1), Eurotiomycetes (n=3), and Chaetothyriomycetes (n=1). The alignment of the amino acid sequences revealed striking differences in homology among protein sequences (data not shown). However, as earlier reported (Fleissner et al., 2005, Craven et al., 2008, Bork & Sudol, 1994), the WW protein-protein interaction domain is highly conserved (70% similarity) within the fungal species analyzed. WW domain contains the motif PPLP and two conserved tryptophan residues spaced 22 amino acid apart. Beyond the WW domain, the Atrophin-1 superfamily and PhoD domains were also identified (Fig. S2B). Comparison of ZtSo protein sequence with its orthologues in the most studied fungi for anastomosis showed 53% identity with *Epichloe festucae;* 54% identity with *Neurospora crassa* and *Sordaria macrospora;* 55% identity with *Fusarium oxysporum*; 60% identity with *Aspergillus oryzae*; and 63% identity with *Alternaria brassicicola*. Phylogenetic analysis grouped the orthologues of the *ZtSo* gene onto three groups based on fungal Classes (Dothideomycetes, Sordariomycetes, and Chaetothyriomycetes together with Eurotiomycetes), independently whether they were parasites, mutualists or saprotrophs (Fig. S2C).

### The mutual attraction towards genetically identical fusion partners is regulated by different genes from MAP kinase pathways

*so*, a gene from the CWI pathway has been shown to have an essential role in self-anastomosis. To determine whether the orthologous gene *ZtSo* plays the same role in *Z. tritici*, we phenotyped the *ΔZtKu70, ΔZtSo*, and *ΔZtSo*-comp strains for the presence of interconnected individuals through anastomoses during growth on WA. Fusion bridges between blastospore germlings or filamentous hyphae were only observed for those strains possessing the *ZtSo* gene. On the other hand, the *ΔZtSo* mutant lost the ability to undergo vegetative fusions (Figs. 3A and B), unveiling the contribution of *ZtSo* for anastomosis between genetically identical *Z. tritici* strains.

**Figure 3.**
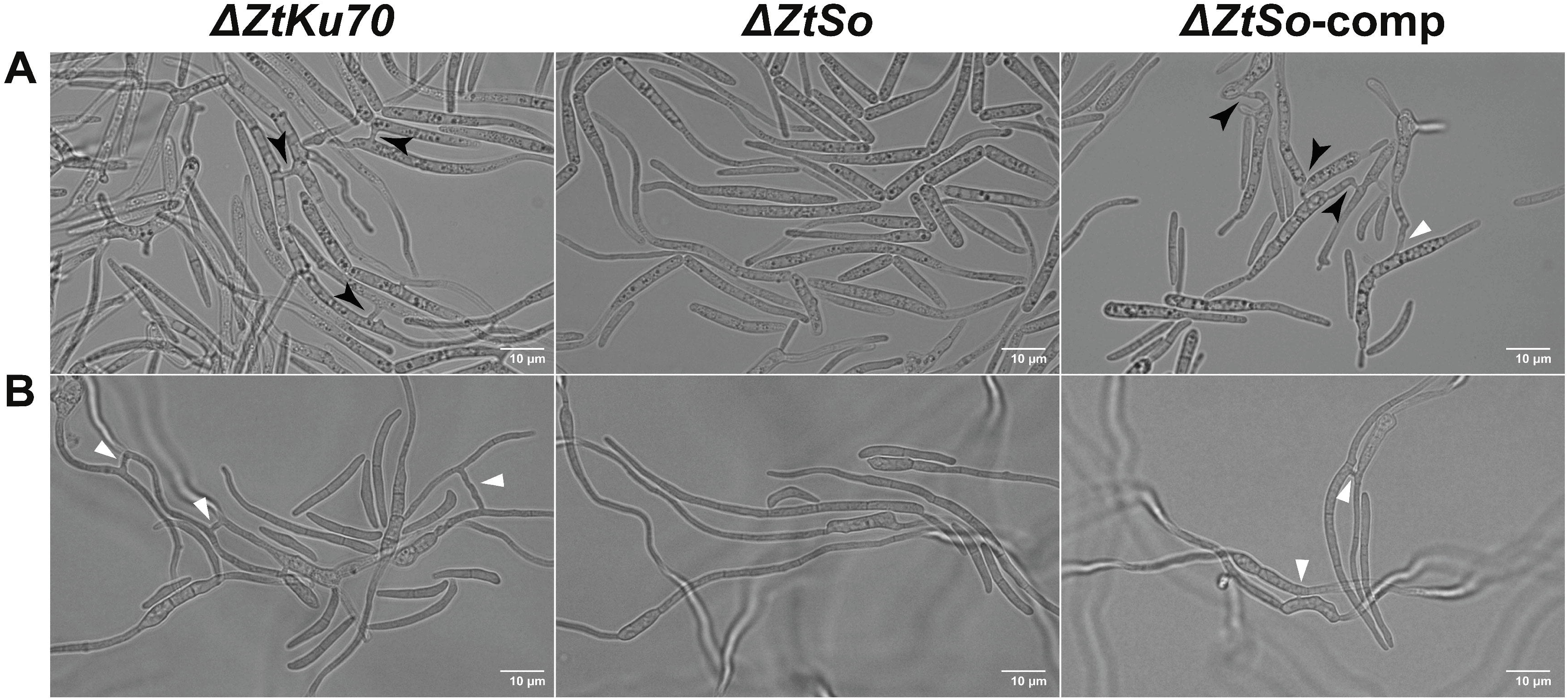
*ZtSo* is required for vegetative cell fusion in *Zymoseptoria tritici*. (A) At initial higher cell density – an inducing condition for CATs, fusion bridges between blastospore germlings were only observed for *ΔZtKu70* and *ΔZtSo*-comp strains. Black arrows indicate the CATs. (B) At initial lower cell density – an inducing condition for VHFs, anastomoses were noticed only for those strain possessing the *ZtSo* gene. CATs or VHFs were never formed between *ΔZtSo* mutant spores. White triangles point to the fusion bridges between two fused hyphal cells.

To ensure the failure of cytoplasmic mixing on those individuals lacking the *ZtSo* gene, we mixed blastospores of each tested strain with 1E4_GFP_ blastospores in a 1:1 ratio and microscopically traced the spores up to 40 hours. Hyphal fusions and the continuous streaming of cytoplasmic green fluorescence coming from the fusion with the 1E4_GFP_ strain were observed for all tested combinations (Figs. S3 and S4), except for the mixture between *ΔZtSo* and 1E4_GFP_ (Fig. S5). This confirmed that *ZtSo* is needed in both fusion partners for a mutual recognition previous to the anastomosis.

To determine whether the MAK-1 and MAK-2 pathways are required for anastomoses in *Z. tritici*, we incubated the knocked-out *ZtSlt2* (orthologous to MAK-1) and *ZtFus3* (orthologous to MAK-2) mutants on WA plates. Neither hyphal attraction nor hyphal fusion was observed for both mutants, however, fusion bridges were frequently found between IPO323 wild-type or *ΔZtSlt2-*complemented fungal cells (Fig. S6). These results indicate the recruitment of both CWI and MAK-2 MAP kinase signaling cascades for the regulation of the self-stimulation and self-fusion in *Z. tritici*.

### Deletion of *ZtSo* has a differential impact on hyphal or blastospore growth

*Z. tritici* alters its growth morphology in detrimental to the nutritional conditions (Francisco *et al.*, 2019). To assess whether vegetative cell fusion affects fungal growth, we determined the radial growth of *ΔZtKu70, ΔZtSo,* and *ΔZtSo*-comp on different culture media. A slightly reduced growth was detected in the *ΔZtSo* colonies compared to those possessing the *ZtSo* gene (Fig. S7A) when it’s grown on WA. On average, the colony radii were 5.31 ± 0.12 for *ΔZtKu70;* 4.87 ± 0.10 for *ΔZtSo;* and 5.50 ± 0.09 for *ΔZtSo*-comp [radial growth (mm) ± standard error] (Fig. S7C). Furthermore, *ΔZtKu70* and *ΔZtSo*-comp formed colonies with highly hyphal dense margins, while *ΔZtSo* exhibited only a few filamentations at the colony periphery (Fig. S7B). No morphological differences were detected between blastospores of the tested strains (Fig. S7D). In contrast, the *ΔZtSo* mutant grew significantly faster than the *ΔZtKu70* and *ΔZtSo*-comp in PDA, a nutrient-rich medium (Fig. S7E). Over time, the relative growth rate of *ΔZtKu70* and *ΔZtSo*-comp were, respectively, 32% and 16% lower than the *ΔZtSo* mutant. Together, these results suggest that the reduced mycelial expansion of the *ΔZtSo* mutant grown in a nutrient-poor medium is a consequence of impairment of VHFs. On the other hand, the *ZtSo* gene may contribute to the blastosporulation and its deletion promotes an imbalance of this mechanism in *Z. tritici*.

### *ZtSo* is required for melanization, but it is dispensable for stress sensitivity

No melanin accumulation was observed in *ΔZtSo* mutant colonies grown *in vitro* (Fig. 4), demonstrating the impact of *ZtSo* deletion for *Z. tritici* pigmentation. Next, we postulated that the *ΔZtSo* mutant could either display cellular integrity defects and/or being susceptible to environmental stresses. We tested nine different abiotic stressors, such as temperature, oxidative, osmotic, cell wall, and cell membrane stresses. Overall, no variability in stress response was noticed among the strains (Fig. 5). However, *ΔZtSo* formed slightly bigger colonies compared to those from *ΔZtKu70* and *ΔZtSo*-comp, most likely to the increase of blastospore formation when its grown- on nutrient-rich media (Fig. S7E). Thus, we found no evidence that the *Zt*S*o* gene is required for the maintenance of the cell wall integrity. Furthermore, our results demonstrated that melanin does not act as “fungal armor” protecting *Z. tritici* against the tested stresses.

**Figure 4.**
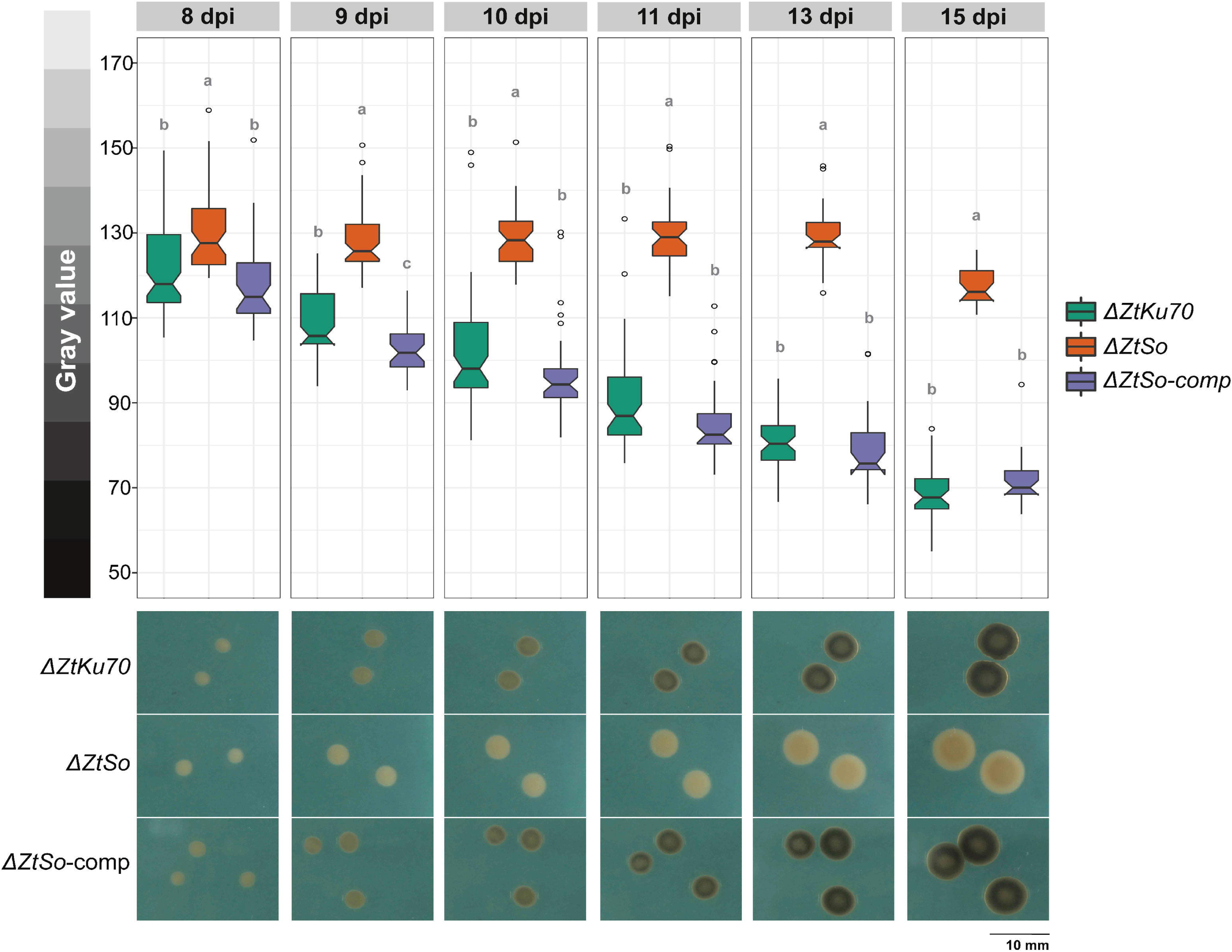
Disruption of *ZtSo* strongly affects the melanization of *Zymoseptoria tritici*. The fusion defective mutant was significantly less melanized than the *ΔZtKu70* and *ΔZtSo*-comp strains, which exhibited higher melanin accumulation over time. Bars represent standard errors of the mean gray value on at least 40 colonies. Different letters on the top of the bars indicate a significant difference among the tested strains according to the Analysis of Variance (ANOVA). The notch displays a 95% confidence interval of the median. Open circles represent the outlier values of each strain. Pictures shown below the bar plot represent the melanization level of *ΔZtKu70, ΔZtSo* and *ΔZtSo*-comp strains. Gray value scale (0 = black and 255 = white) is shown on the left.

**Figure 5.**
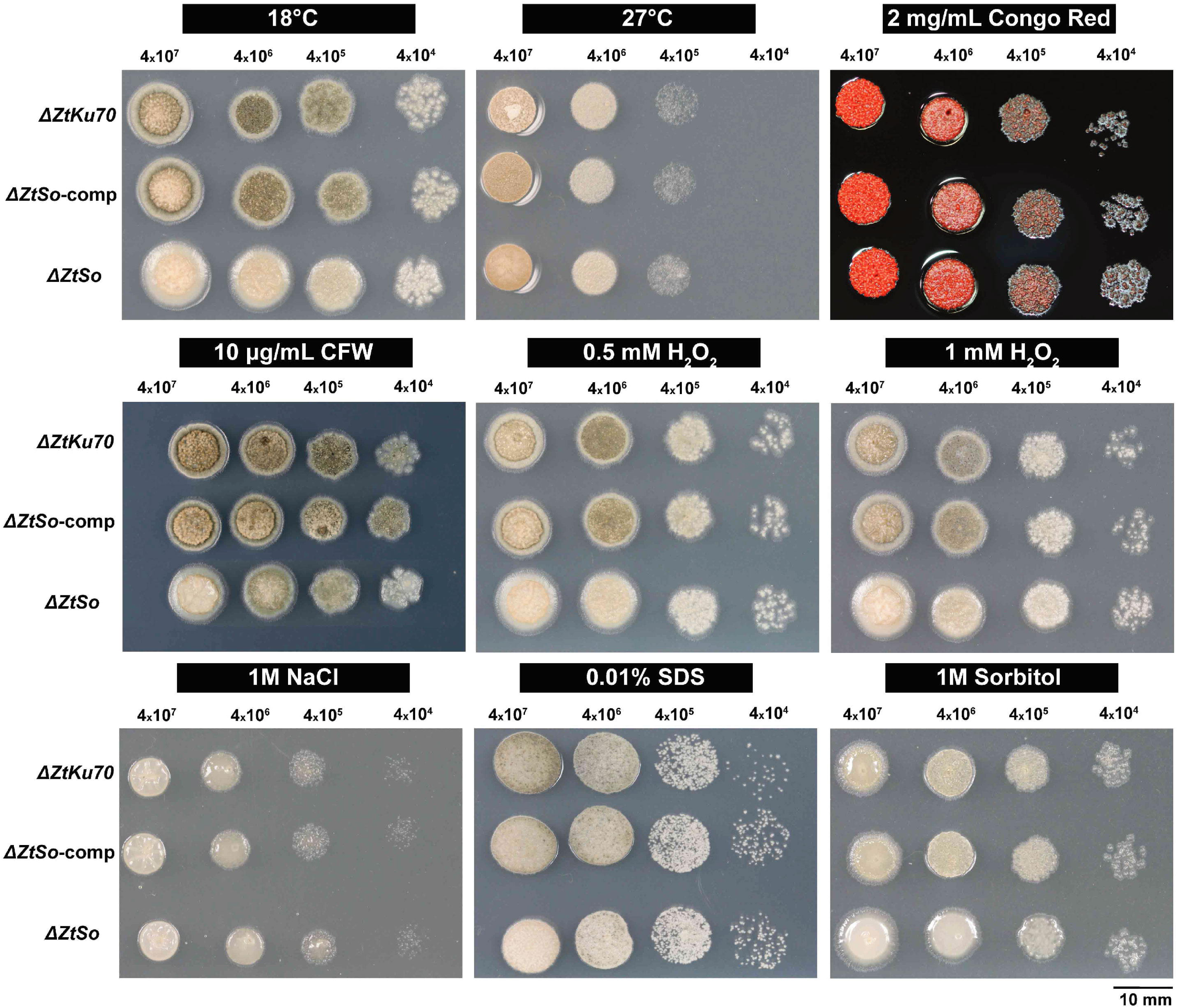
*ZtSo* is dispensable for the cellular integrity. A serial dilution of blastospore suspension from *ΔZtKu70, ΔZtSo,* and *ΔZtSo*-comp strains were exposed to nine different stress conditions, including different temperatures (18°C and 27°C), oxidative stress (0.5 and 1 mM of hydrogen peroxide - H_2_O_2_), osmotic stress (1M sodium chloride - NaCl and 1M sorbitol), cell wall stress (2 mg/mL congo red and 10 μg/mL calcofluor white - CFW), and plasma membrane stress (0.01% sodium dodecyl sulfate-SDS) for five days. Besides the impairment in melanization exhibited by the *ΔZtSo* mutant, all tested strains do not vary in their tolerance to different cellular stressor agents.

### Hyphal fusions are essential for the development of asexual fruiting bodies

We inoculated a susceptible wheat cultivar with the tested *Z. tritici* strains to assess the biological role of vegetative cell fusion during the pathogen lifecycle *in planta*. Typical symptoms caused by *Z. tritici* infections were visible after 9 days post-infection (dpi) and the disease progression was similar among the plants inoculated with *ΔZtKu70, ΔZtSo* or *ΔZtSo*-comp strains (Figs. 6A and S8). This indicates that *ZtSo* is neither required for host penetration nor the asymptomatic or necrotrophic phases of the fungus. The asexual fruiting bodies (pycnidia) were visible on plants inoculated with those strains possessing the *ZtSo* gene at 14 days post-inoculation (dpi). In contrast, plants infected with *ΔZtSo* never developed pycnidia. The failure to form the asexual reproductive structure was also observed in other susceptible cultivars to *Z. tritici* (Fig. S9).

**Figure 6.**
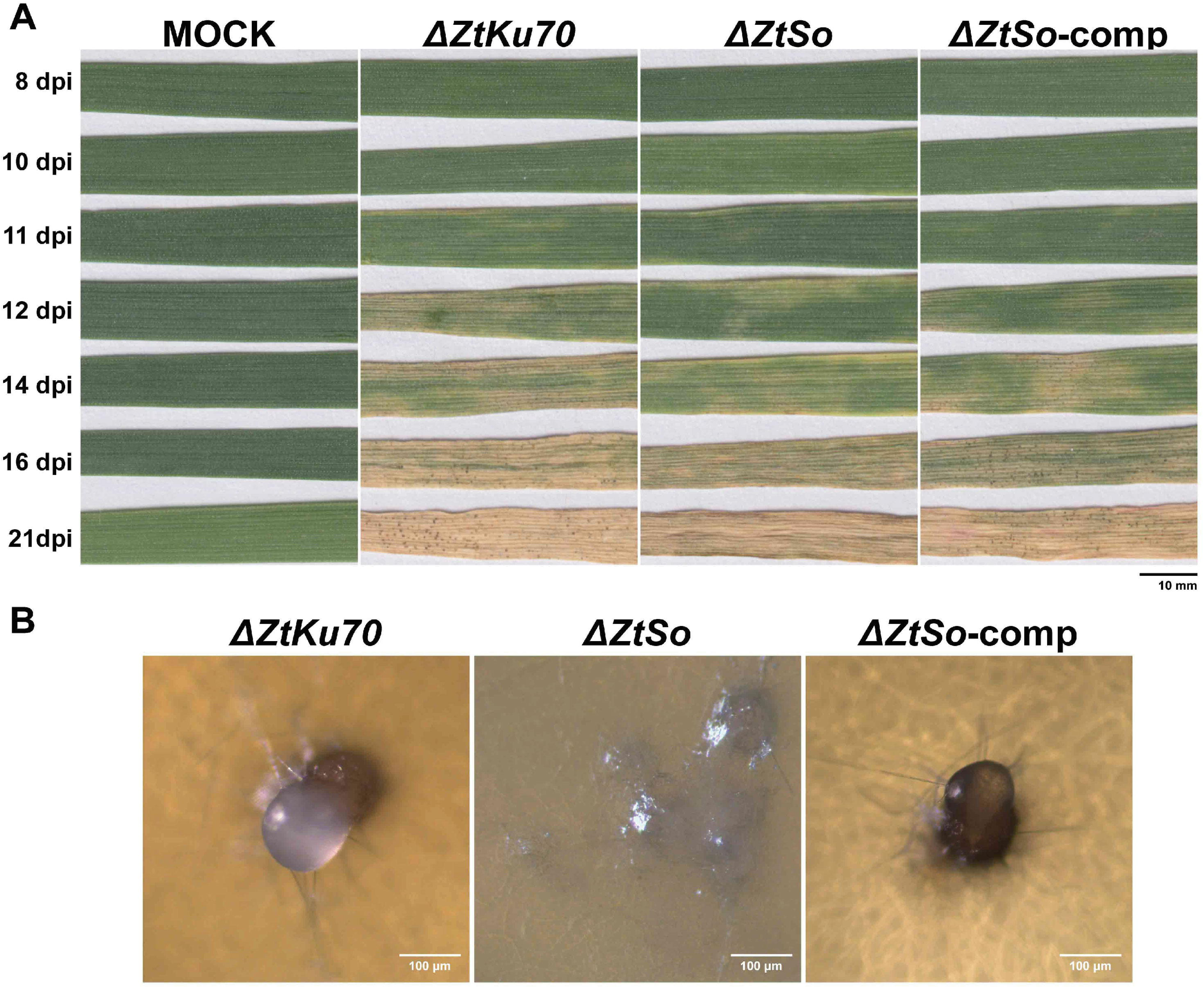
The biological role of vegetative hyphal fusions for the disease progress and development of the asexual reproductive structures of *Zymoseptoria tritici*. (A) Susceptible wheat cv. Drifter inoculated with *ΔZtKu70, ΔZtSo,* or *ΔZtSo*-comp strains were evaluated up to 21 days post-infection (dpi). All tested strains exhibited similar disease progress. The onset of the necrotrophic phase started at 11 dpi, whereas the necrotic lesions continued to expand across the leaves over time, resulting in the collapse of the host tissue at 21 dpi. Pre-pycnidia were noticed at 12 dpi, and the mature pycnidia are prominent after 14 days of incubation. However, neither pre-pycnidia nor mature pycnidia were formed in the wheat plants inoculated with the fusion defective *ΔZtSo* mutant. (B) Blastospores of *ΔZtKu70, ΔZtSo,* and *ΔZtSo*-comp strains were inoculated on wheat extract agar plates at 18°C and incubated under UV-A light (16:8 light:dark cycle). After 20 days of incubation, *ΔZtKu70* and *ΔZtSo*-comp strains produced brown pycnidia-like structures exuding a whitish liquid similar to the oozed cirrhus-containing pycnidiospores spores observed for *Z. tritici*-infected wheat plants. In contrast, *ΔZtSo* mutant formed only mycelial knots, which are aggregates of filamentous hyphae and the precursor developmental stage of pycnidium, but the mature asexual fruiting bodies were never formed.

To distinguish whether the absence of pycnidia formation would be either a consequence of the lack of hyphal fusions or higher sensitivity of the *ΔZtSo* mutant to the plant defenses, we used a wheat extract agar medium to induce pycnidia formation *in vitro. ΔZtSo* produced only mycelial knots, the forerunner developmental stage of mature pycnidia, but no asexual reproductive structures were further developed. Unlike, the pycnidia-like structures formed by *ΔZtKu70* or *ΔZtSo*-comp strains were exuding a whitish liquid similar to the oozed cirrhus-containing pycnidiospores observed *in planta* (Fig. 6B).

Therefore, we next postulated that *ΔZtSo* has a similar colonization pattern than the wild-type strain, including accumulation of hyphae in the sub-stomatal cavity, but the lack of cell fusion would obstruct the pycnidial development. We monitored the wheat plants infected with 1E4_GFP_ or 1E4_GFP_*ΔZtSo* strains using confocal microscopy up to 12 dpi. At the earlier evaluated stages of the plant infection, we observed mainly host penetration, initial intercellular hyphal extension and sub-stomatal colonization (Fig. S10 – 6 and 7 dpi). None difference was noticed in fungal development, however, hyphal fusions established during epiphytic host colonization were only found for 1E4_GFP_ strain. At 8 and 9 dpi, we observed that the first intercellular hyphae surrounding the stomatal guard cells produced specialized knots from where secondary hyphae emerge and germinate (Figs. 7 and S10). These secondary hyphae fuse with other nearby hyphae, creating an interconnected network in the sub-stomatal cavity (e.g. for the 1E4_GFP_ strain) or it keeps extending as individual hypha (e.g. for the 1E4_GFP_*ΔZtSo* mutant) (Figs. 7 and S10). The combination of sub-stomatal hyphal accumulation and anastomoses generates the mature pycnidium, which later supports the asexual reproduction of *Z. tritici*. On the other hand, the lack of anastomosis stops the development of the pycnidium and, consequently, impair the asexual cycle of the fungus. Thus, we concluded that hyphal fusions are crucial for the pycnidial development and the disturbance of this mechanism ceases the asexual reproduction of *Z. tritici*.

**Figure 7.**
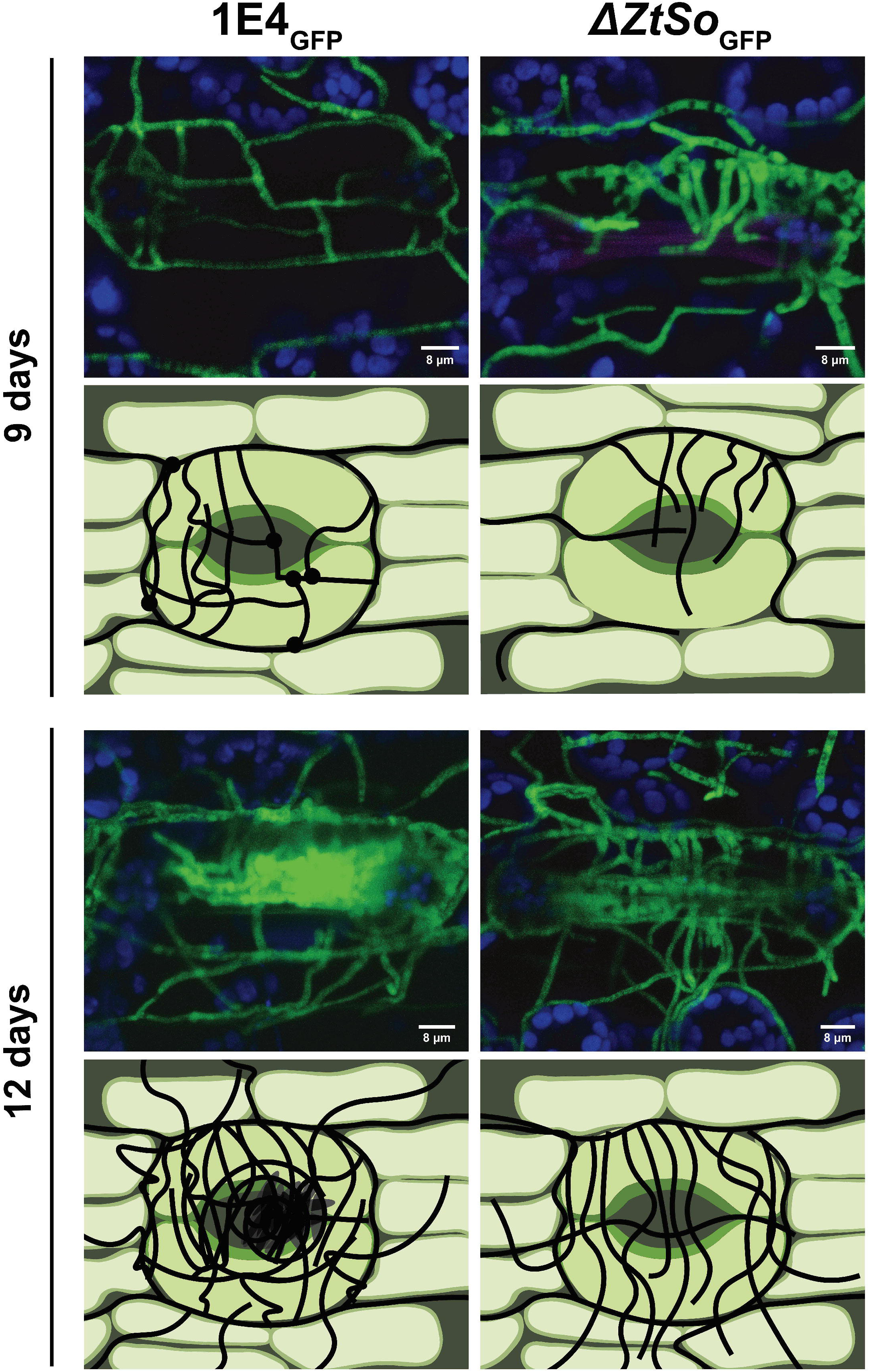
Schematic demonstration of pycnidial development during plant infection by a filamentous fungus. Susceptible wheat cv. Drifter was inoculated with the fluorescent *Zymoseptoria tritici* 1E4_GFP_ (wild-type) and *ΔZtSo*_GFP_ strains and monitored by confocal microscopy at different days post-infection (dpi). For earlier time points (6, 7 and 8 dpi), please see Fig.S10. The first intracellular hyphae surrounding the stomatal guard cells produced specialized knots from where secondary hyphae emerge and germinate. After 9 days of infection, these secondary hyphae from the 1E4_GFP_ strain fuse with another nearby hypha (represented by black circles), creating an interconnected network in the sub-stomatal cavity. The combination of sub-stomatal hyphal accumulation and anastomoses generates the pre-pycnidium at 12 days, which later supports the asexual reproduction of *Z. tritici.* Unlike, the filamentous hyphae from the *ΔZtSo*_GFP_ mutant kept extending as individual hyphae and none fusion point was observed. The lack of anastomosis stops the developmental process of the pycnidium and, consequently, impair the asexual cycle of *Z. tritici*.

## Discussion

Cell-to-cell communication regulates a myriad of biological process that drive fungal development and ecological diversifications. The sophisticated evolution in the fungal language allowed a fine-tune coordination of signal senders and receivers and, consequently, the regulation of complex signaling networks. Here, we explored the functional relationship of a gene involved in cell-to-cell communication and its biological contribution to the development and fitness of a filamentous fungus plant pathogen.

Vegetative cell fusion is one of the most important cellular developmental processes of a mycelial fungal colony (Glass et al., 2000, Simonin *et al.*, 2012). The cytoplasmic continuity generated by cell fusion provides adaptative advantages to the interconnected mycelial network essentially for resource sharing and introgression of genetic material (Simonin et al., 2012, Roca *et al.*, 2005b, Mehrabi et al., 2011). CATs forming at earlier stages of the vegetative growth have been documented for several filamentous fungi (Roca et al., 2005a). We observed that these specialized fusion bridges require a certain cell density, indicating that *Z. tritici* may induce CATs at a critical concentration of a self-produced extracellular molecule, most likely a quorum-sensing molecule. A CAT inducer signal based on a quorum sensing was previously proposed for *N. crassa* and *Venturia inaequalis* (Roca et al., 2005b, Read *et al.*, 2012), but the nature of this molecule and its receptor on the cell remain unknown. On the other hand, the induction of VHF in *Z. tritici* does not seem to be associated with a quorum-sensing because anastomoses were frequently observed at lower initial cell concentration and thus, VHF may be induced by some other environmental signal. For instance, VHFs were pronounced in nutrient-limited conditions in *Z. tritici*, as observed for the causal agent of anthracnose disease, *Colletotrichum lindemuthianum* (Ishikawa et al., 2010). Plant pathogens typically experience nutrient limitations while growing on leaves and the perception of a nutrient-limited environment may act as a stimulus to induce cell fusion in foliar plant pathogens. Both germinating blastospores or pycnidiospores of *Z. tritici* undergo CAT and VHF *in vitro* and *in planta*, supporting our previous observation (Francisco et al., 2019). Thus, we demonstrated that vegetative cell fusions occur independently of the morphotype and it may have an important contribution to the lifestyle of *Z. tritici* during the plant-pathogen interaction.

Unlike the majority of fungal species that form a multinuclear hyphal network (Roper *et al.*, 2011), *Z. tritici* has only one nucleus per septal compartment (Kilaru *et al.*, 2017). We used a GFP-tagged nucleus strain, to track the nuclei movements between two *Z. tritici* fused partners. Nuclei transfer between two encountering hyphae has been described for different fungal species (Chagnon, 2014, Mehrabi et al., 2011, Roper et al., 2011). The consequences of genetic exchange include the formation of viable heterokaryons (Roper et al., 2011) and the risk to introduce pathogenic elements or virulence genes (Biella et al., 2002, Goddard & Burt, 1999, Friesen et al., 2006). This raises the question of whether heterokaryon formation could happen in *Z. tritici* after cell fusions. We showed that after the fusion bridge formation connecting the two identical *Z. tritici* cells, the new cell compartment is occupied by a migrating nucleus coming from a neighboring nucleus-divided by mitosis. Heterokaryon cells were never observed neither near nor far from the anastomosis point, indicating that *Z. tritici* may have evolved to limit the spreading of genetic elements and restricting the formation of heterokaryons and the parasexual cycle. However, we do not discard the possibility that exchanges of DNA fragments or pathogenic elements may affect the fitness and evolution of this fungus. For instance, transposable elements (TEs) are extraordinary generators of fungal diversity and versatility (Mat Razali *et al.*, 2019, Castanera *et al.*, 2016). These selfish elements, in particular, the DNA Transposon (or Class II TE), that represents 14.6% of the repetitive DNA in *Z. tritici* (Dhillon *et al.*, 2014) and uses the transposase activity to *“cut and paste”* genome sequences, could mediate gene or chromosome horizontal transfers. Notwithstanding, this mechanism was proposed for the *ToxA* neighboring type II hAT-like transposase gene horizontally transferred between three wheat pathogen-related species (McDonald *et al.*, 2019). Likewise, the mycoviruses existing within the cytoplasm can also be horizontally transmitted during anastomosis (Ihrmark *et al.*, 2002, Pearson *et al.*, 2009). *Z. tritici* is known to host double-stranded RNA (dsRNA) mycoviruses species (Zelikovitch *et al.*, 1990, Kema *et al.*, 2008), including a homologous dsRNA hypovirus identified in *Fusarium graminearum* and associated with altered fungal growth, pigmentation and reduced virulence (Chu *et al.*, 2002). Hypovirulence associated with dsRNA mycoviruses has been reported for several plant pathogen fungi (Nuss, 2005), but the impact of these obligate parasites on *Z. tritici* biology remain broadly unknown. Here, we propose that cell fusions may generate non-conventional possibilities to contribute to the high genetic diversity observed in *Z. tritici* isolates. Whether these genetic elements or mycoviruses can be acquired by anastomosis and impact *Z. tritici* diversification will be a subject of futures studies.

In the last decades, the identification of fusion defective mutants contributed to the understanding of the molecular mechanisms underlying cell communication and fusion, especially by unveiling the interplay of the mitogen-activated protein (MAP) kinase pathways during chemotropic interaction (Fleissner et al., 2009). MAP kinase pathways are involved in extracellular signal perception and regulation of diverse genes essential for mating, filamentation, pathogenicity, cell integrity and stress response (Deng *et al.*, 2018, Zhao *et al.*, 2007, Pandey et al., 2004, Maddi *et al.*, 2012, Leng & Zhong, 2015, Hagiwara *et al.*, 2016) and thus, the deletion of MAP kinase-related genes result in pleiotropic phenotypes due to the interaction in multiple biological processes. Though fungal communication and fusion require the regulation of several genes (Fischer & Glass, 2019), the crosstalk between cell wall integrity (CWI) and MAK-2 signal response, two conserved MAP kinase signaling pathways, are essential to produce, secrete and sense the chemoattractant molecule produced during cell fusion (Fleissner et al., 2009). The deletion of MAK-1 or MAK-2 genes disrupts the signaling cascade affecting the self-anastomosis in filamentous fungi. We used the *ZtSlt2* (MAK-1) and *ZtFus3* (MAK-2) *Z. tritici* orthologues to demonstrate the functional conservation of these MAP kinase pathways in this fungus. Hitherto, *ZtSlt2* and *ZtFus3* were known as essential genes for the pathogenicity of *Z. tritici,* regulating invasive growth and host penetration (Mehrabi *et al.*, 2006, Cousin *et al.*, 2006). Here, we showed that both *ΔZtSlt2* and *ΔZtFus3* lost their self-stimulation and were unable to undergo anastomosis. Our results indicate that cell fusions in *Z. tritici* follow the ping-pong self-signaling mechanism described for *N. crassa* (Fleissner et al., 2009, Read et al., 2009), where the signal sending and receiving is coordinated by genes-associated with the CWI and MAK-2 pathways. Besides, this is the first report of CWI and MAK-2 pathways regulating cell communication in *Z. tritici*.

To evaluate the impact of vegetative cell fusion for the biology of a latent necrotrophic fungus, we used *ZtSo* orthologous of *N. crassa so*. In contrast to the MAP kinase-related genes, the characterization of *so* orthologues results in less pleiotropic phenotypes (Fleissner et al., 2005, Prados Rosales & Di Pietro, 2008, Craven et al., 2008). Filamentous fungus lacking *so* gene is impaired in self-anastomosis (Craven et al., 2008, Prados Rosales & Di Pietro, 2008, Charlton et al., 2012, Fleissner et al., 2005), including *Z. tritici*, in which we showed that the deletion of *ZtSo* abolishes the vegetative cell fusion of this fungus. We characterized the impact of this fusion defect during different developmental stages of *Z. triti*ci. For example, while the fusion competent individuals have an advantageous growth on WA medium promoting larger colonies with dense hyphal borders, the *ΔZtSo* mutant exhibited an asymmetrical hyphal growth extension. It has been previously demonstrated that the direction of nutrient distribution occurs mostly from the central part of mycelium to outwards and its streaming speed is driven by the anastomosis (Simonin et al., 2012). Thus, the repression of colony extension observed for the *ΔZtSo* mutant grown on a nutrient-limited environment may be a consequence of the irregular distribution of cytoplasmic content, including this limited-food resource, throughout the mycelial colony. Surprisingly, *ΔZtSo* underwent extensive production of blastospores, generating larger yeast-like colonies than the fusion competent individuals in a nutrient-rich environment. This result suggests that the increased growth rate observed for the fusion-defective mutant may reflect the role of the *ZtSo* gene on the vegetative growth, in this case, the boost of the blastosporulation mechanism, probably due to an interplay of different signaling pathways. The genetic relationship between the derepression of blastosporulation and the lack of *ZtSo* gene remains to be elucidated.

We showed that the *ΔZtSo* does not accumulate melanin, resulting in whiter and larger colonies than those formed by *ΔZtKu70* and *ΔZtSo*-comp strains. These findings illustrate how melanization drove pathogen adaptation to a trade-off between energy cost for pigment production and fungal growth. The deleterious effect on *Z. tritici* growth caused by a higher accumulation of melanin was previously reported for this fungus (Krishnan *et al.*, 2018). Melanins are dark-pigmented secondary metabolites often associated with the fungal cell walls. Though fungi can produce different kinds of melanins, hitherto, it has been suggested that melanization of *Z. tritici* is controlled only by the polyketide synthase (PKS) gene cluster containing catalytic enzymes and transcription regulators of the 1,8-dihydroxynaphthalene (DHN) melanin (Lendenmann *et al.*, 2014, Krishnan et al., 2018). The CWI pathway, to which *ZtSo* belongs, is the central signaling cascade regulating diverse biological processes, including the production of secondary metabolites (Levin, 2005, Valiante, 2017, Park *et al.*, 2008). It has been demonstrated to plant-pathogenic fungi that the deletion of CWI-associated genes inhibited pigmentation by reducing the expression of DHN melanin biosynthetic genes (Yago et al., 2011, Liu et al., 2011, Valiante et al., 2015). Thus, we speculate that the CWI signaling pathway regulates the PKS-encoding genes leading to the melanin accumulation in *Z. tritici*. The deletion of *ZtSo* may impair the CWI-regulatory cascade and, consequently, the regulation of DHN-melanin production, resulting in the lack of pigmentation observed for both *ΔZtSo* and *ΔZtSlt2* (Mehrabi et al., 2006) mutants from the CWI pathway. On the other hand, *so* is also described for its putative function in the secretion of internal vesicles transporting the chemoattractant molecule at the growing cell tip (Fleissner et al., 2005, Fleissner & Herzog, 2016). Considering that different studies have demonstrated that fungal melanin may be synthesized in internal vesicles and transported to the cell wall (Rodrigues *et al.*, 2008, Silva *et al.*, 2014), we propose an alternative hypothesis to explain the impaired melanization by *ΔZtSo* mutant. The disruption of *ZtSo* may affect the transport of internal vesicles carrying the pigment to the cell wall or the laccase oxidase enzyme required to polymerize 1,8-DHN to form the DHN-melanin polymer (Plonka & Grabacka, 2006), resulting in a non-melanized strain. Both hypotheses require further investigations.

Melanin has also been postulated to contribute to fungal protection against fungicide and environmental stresses in *Z. tritici* (Krishnan et al., 2018, Lendenmann et al., 2014) and its regulation depends on environmental cues and colony development (Lendenmann et al., 2014), though not yet completely understood. We used nine different cellular stressors to evaluate whether (i) the defect in melanin accumulation or (ii) the deletion of ZtSo, the scaffold protein for the MAP kinase genes from the CWI pathway, would affect pathogen stress tolerance. Since the non-melanized *ΔZtSo* mutant displayed the same degree of stress sensitivity than *ΔZtKu70* and *ΔZtSo*-comp strains, we concluded that ZtSo does not act as a scaffold protein for all CWI pathway function. This observation was previously described for the model fungus *Sordaria macrospora*, where the PRO40 (orthologous to ZtSo) operates as a scaffold for the CWI-encoding genes during fungal development, hyphal fusion, and stress response, but not for growth under cell-wall stress agents (Teichert *et al.*, 2014). Our results showed that melanin accumulation of *Z. tritici* does not promote an advantage of fungal survival in harsh environments, at least for the stressful conditions tested in this study.

We demonstrated that VHFs are dispensable for the pathogenicity of *Z. tritici*. The fusion-defective *ΔZtSo* mutant displayed a similar host damage progression than those individuals possessing the gene. This finding exemplifies the distinct effects of cell fusions on fungal pathogenicity. For instance, for the soil-borne *Fusarium oxysporum*, VHF-impaired mutants exhibited only a slightly reduced virulence, whereas, for the necrotrophic plant pathogen *A. alternata,* VHFs are necessary for the full virulence of the fungus (Prados Rosales & Di Pietro, 2008, Craven et al., 2008). Though the deletion of *ZtSo* is not essential for host penetration, colonization or for the onset of the necrotrophic phase *per se*, it is during hyphal accumulation in the sub-stomatal cavity that the fusion defect impacts *Z. tritici* fitness. As far as we know, this is the first time that pycnidial development has been microscopically detailed in a filamentous fungus. We showed that the first intercellular hyphae surrounding the stomatal guard cells produced specialized knots from where secondary hyphae emerge and germinate to fuse with other adjacent hyphae. Consequently, this preliminary hyphal network creates the basis for a symphogenous development that builds the concave-shaped pycnidial wall of the mature pycnidium. On the other hand, the inability to undergo anastomosis ceases the development of the asexual fruiting bodies of the fungus, and therefore, abolish fungal reproduction. The accumulation of hyphae observed in the sub-stomata chamber by the *ΔZtSo* mutant confirms that there is a specific signal that triggers sub-stomatal hyphal aggregation, which is independent of the chemoattractant molecule secreted by the fungus to induce hyphal fusion. Regarding the sexual phase, we did not investigate the impact of the *ΔZtSo* for the sexual cycle, because is not yet feasible to generate *in vitro* crosses for *Z. tritici*. Fleissner et al. (2005) demonstrated that the deletion of *so* in *N. crassa* affects female fertilization, which blocks the sexual reproduction of the mutant. Further experiments need to be performed to address this question for *Z. tritici*, but we believe that the *ZtSo* gene would also play a crucial role in the sexual reproduction of this pathogen.

In summary, the characterization of *ZtSo* gene demonstrated its fundamental role in fungal biology. Beyond the impact of *ZtSo* for self-stimulation and self-fusion, we show the contribution of this gene for fungal development, including a negative impact on hyphal extension, derepressing of blastosporulation and impairment in melanization. Besides, we demonstrated that VHFs are dispensable for pathogenicity, but essential for pycnidial development. Taken together, our data illustrate how cell fusion affects *Z. tritici* fitness and provide a powerful new gene target to control Septoria tritici blotch (STB) disease.

## Experimental Procedures

### Strains and growth conditions

The Swiss *Z. tritici* strain ST99CH_1E4 (abbreviated as 1E4), described by Zhan *et al.* (2002), and mutant lines derived from this strain, as well as the 1E4 strain expressing cytoplasmic GFP (1E4_GFP_) or mCherry (1E4_mCh_), were used in this study. The knocked-out *ΔZtSlt2* (Mehrabi et al., 2006) and *ΔZtFus3* (Cousin et al., 2006) mutants were provided by Marc-Henri Lebrun (National Institute of Agricultural Research – INRA, France).Because the MAP kinase mutants were generated in the genetic background of IPO323 (Kema & van Silfhout, 1997), this strain was also used. Routinely, *Z. tritici* was cultivated on yeast-sucrose broth (YSB) medium (10 g/L yeast extract, 10 g/L sucrose, 50 μg/mL kanamycin sulfate; pH 6.8). Each strain was stored in glycerol at −80°C until required and then recovered in YSB medium incubated at 18°C for four days.

### Plant infection to obtain fluorescent pycnidiospores

Wheat seedlings from the susceptible wheat cultivar Drifter were grown for 16 days in the greenhouse at 18°C (day) and 15°C (night) with a 16h photoperiod and 70% humidity. Blastospore suspensions of 1E4_GFP_ or 1E4_mCh_ were obtained after four days of growth in YSB medium. Spore suspensions were adjusted to a final concentration of 10^6^ blastospores/mL in 30 mL of sterile water supplemented with 0.1% (v/v) Tween and applied to run-off using a sprayer and the plants were kept for three days in sealed plastic bags, followed by 21 days in a greenhouse. Leaves with pycnidia were harvested and transferred to a 50 ml Falcon tube containing sterile water and gently agitated to harvest the pycnidiospores. Fluorescent pycnidiospores were used to assess the cell fusion events during *in vitro* and *in vivo* growth. Pycnidiospore suspension-tagged GFP or mCherry were also adjusted to a final concentration of 10^6^ pycnidiospores/mL and a new batch of plants were inoculated as described above. Plants co-infected by both 1E4_GFP_ and 1E4_mCh_ strains were used to observe VHFs on the wheat leaf surface.

### Characterization of cell fusion events *in vitro* and *in vivo*

The ability of *Z. tritici* to undergo cell fusions was evaluated using blastospores and pycnidiospores of 1E4_GFP_ and 1E4_mCh_. Cell concentrations were adjusted to 3.3×10^7^ blastospores/mL or 3.3×10^6^ blastospores/mL to induce CATs or VHFs, respectively. 300 μL of each morphotype and fluorescence was plated on water agar (WA) to create a ratio of 1:1 and to provide a final concentration of 10^7^ blastospores/mL or 10^6^ blastospores/mL. A section of about 1 cm^2^ of agar was aseptically cut and placed on a microscope slide. The mixing of both cytoplasms content through CATs or VHFs was checked up to 40 hai using a Leica DM2500 fluorescent microscope with LAS v.4.6.0 software. GFP excitation and emission was at 480/40 nm and 527/30 nm, respectively, whereas, mCherry was excited at 580/20 nm and detected at 632/60 nm.

VHFs during spore germination on wheat leaf surface were obtained via confocal images using a Zeiss LSM 780 inverted laser-scanning microscope with ZEN Black 2012 software. An argon laser at 500 nm was used to excite GFP fluorescence and chloroplast autofluorescence, while mCherry excitation was at 588 nm. The emission wavelength was 490-535, 624-682, and 590-610 nm for GFP, chloroplast autofluorescence, and mCherry, respectively. Plants co-inoculated with blastospores or pycnidiospores of 1E4_GFP_ and 1E4_mCh_ strains were checked up to 48 hai.

### Genetic exchange during anastomosis

To determine whether nuclei exchange happens during VHFs in *Z. tritici*, we used the IPO323 ZtHis1-ZtGFP strain (Kilaru et al., 2017), which has the GFP as a fluorescent marker labeling the nucleus. Blastospores of the IPO323 ZtHis1-ZtGFP strain were plated on WA plates at a final concentration of 10^6^ blastospores/mL. Spores were monitored up to 72 hai by light and fluorescent microscope.

### Orthologues identification, protein alignment and phylogenetic analysis

The *so* gene sequence from *Neurospora crassa* (XM_958983.3) was blasted against the *Z. tritici* genome (https://genome.jgi.doe.gov/Mycgr3/Mycgr3.home.html) to identify its orthologous in this fungus. The *Z. tritici* So orthologous protein sequence was used for a Blastp analysis against the NCBI database (National Center of Biotechnology Information). Blastp searches at expected value homology cut-off of 1e-10 were included as positive. A dataset containing So orthologues proteins of different Ascomycete species were used for phylogenetic analysis. Three members of Basidiomycetes were used as outgroup. Protein sequences were aligned using AliView program (Larsson, 2014). The best-fit model of amino acid evolution was the LG+G, determined by Mega6 software (Tamura *et al.*, 2013). Amino acid sequences were aligned using Muscle followed by maximum likelihood phylogeny reconstruction using 1,000 bootstraps and performed with the software Mega6 (Tamura et al., 2013).

### Plasmid constructions and transformations

All PCRs for cloning procedures were performed using NEB Phusion polymerase (New England Biolabs). Primers used for cloning, sequencing and knock-out confirmations are listed in Supporting Information Table S1. DNA assemblies were conducted with the In-Fusion HD Cloning Kit (Takara BIO) following the manufacturer’s instructions. To increase the homologous recombination efficiency, we first inactivated the *ZtKu70* (*Mycgr3G85040* or *Zt09_3_00215*) gene in the1E4 strain using the plasmid pGEN-YR-*ΔZtKu70* (Sidhu et al., 2015), containing a geneticin resistance gene cassette (also known as G418), as a selectable marker. To disrupt the *Z. tritici so* gene (*ZtSo*), 1 Kb size of both flanking regions were amplified from the 1E4 genomic DNA. The hygromycin resistance gene cassette (*hph*), used as a selective marker, was amplified from pES6 plasmid (obtained from E. H. Stukenbrock, Kiel University, unpublished). The pES1 plasmid (obtained from E. H. Stukenbrock, Kiel University, unpublished) was digested with *Kpn*I and *Sbf*I for plasmid linearization, and three fragments were assembled, which resulted in the pES1-*ΔZtSo.* To reintroduce the *ZtSo* gene into the *ΔZtSo* mutant strain, we used the plasmid pES1. We amplified the nourseothricin resistance gene cassette (*nat*) from pES43 plasmid (obtained from E. H. Stukenbrock, Kiel University, unpublished) to be used as a selectable marker. *ZtSo* gene containing 1 Kb size of each flank region and the *nat* resistance gene cassette was assembled into pES1, resulting in pES1-*ΔZtSo*-comp that allowed to introduce the *ZtSo* gene into its native location (Fig. S11). We also used the pES1-*ΔZtSo* plasmid to knock-out the *ZtSo* gene in the 1E4_GFP_ genome background, enabling the visualization of the GFP-tagged mutant during host infection.

Plasmids were transformed into *E. coli* NEB 5-alpha by heat shock transformation for plasmid propagations, followed by plasmid miniprep using QIAprep Spin Miniprep (Qiagen) according to manufacturer’s instructions. Successful plasmid constructions were confirmed by Sanger sequencing before been transformed into *Agrobacterium tumefaciens* strain AGL1 cells by electroporation.

*Z. tritici* 1E4 strain was transformed by *Agrobacterium tumefaciens*-mediated transformation (ATMT) according to Meile *et al.* (2018). The knock-out of the target genes was verified by a PCR-based approach using a forward primer specific to the upstream sequence of the disrupted gene and a reverse primer specific to bind in the resistance cassette (Table S1). We determined the copy number of the transgene by quantitative PCR (qPCR) on genomic DNA extracted with the DNeasy Plant Mini Kit (Qiagen). We used as qPCR target gene the selection marker and the *TFIIIC1* or *18S*, as reference genes (Table S1). Lines with a single insertion were selected for further experiments.

### Phenotypic characterizations

For all phenotypic analyses, *ΔZtKu70* was considered the wild-type (WT) strain. To pinpoint the role of the *ZtSo* gene on the vegetative cell fusion, blastospore suspension of *ΔZtKu70, ΔZtSo,* and *ΔZtSo*-comp was added to a final concentration of 10^6^ blastospores/mL or 10^7^ blastospores/mL into WA and incubated at 18°C to induce CATs or VHFs, respectively. For the MAP kinase *ΔSlt2* and *ΔFus3*, IPO323 and *ΔSlt2*-complemented strains, WA plates were inoculated only at a final concentration of 10^6^ blastospores/mL. Cell fusion events were monitored up to 40 hai by light microscopy. Because fusion bridges were not observed between individuals lacking *ZtSo*, we mixed 150 μL of 10^6^ blastospores/mL of *ΔZtKu70, ΔZtSo,* or *ΔZtSo*-comp strains with the same concentration of 1E4_GFP_ blastospores in a ratio of 1:1 to confirm the failure of cytoplasm streaming. At least 50 spores of each sample combination were monitored by light and fluorescent microscopy..

To test for altered fungal growth, we used PDA (39 g/L potato dextrose agar, 50 μg/mL kanamycin sulfate) and WA media to induce blastospore and hyphal growth, respectively. 200 μL of spore suspension of *ΔZtKu70*, *ΔZtSo*, and *ΔZtSo*-comp were plated at a final concentration of 2×10^2^ blastospores/mL on each aforementioned media and incubated at 18°C. At least 40 colonies formed in five independent PDA plates were photographed from the bottom using a standardized camera setting (Lendenmann et al., 2014) at 8, 9, 10, 11, 13, and 15 days post-incubation (dpi). Digital images were processed using a macro developed in the ImageJ software (Schneider *et al.*, 2012), which scores the area of individual colonies in the images. Fungal growth was obtained by converting the colony area into radial growth (millimeter (mm)) based on the formula 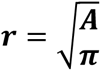, for each strain. Radial growth values were plotted in a boxplot graphic using the ggplot2 package from R (Wickham, 2009). Analysis of variance (ANOVA) was performed to determine the differences in fungal growth among the strains using the agricolae package in R (Mendiburu, 2015). The radial growth rate (mm/day) for each strain was measured by plotting the colony radius over time, which fitted to a linear model (Pearson’s correlation coefficient value (r^2^ ≥ 0.98)). Relative growth rate was calculated by dividing the slope of the regression line of *ΔZtSo* by the slope of *ΔZtKu70* or *ΔZtSo*-comp strains.

Because mycelial growth on the WA plate exhibits a poor color contrast, it was not possible to use the macro to estimate mycelial area. Thus, we manually calculated the mycelial diameter from digital images of at least 40 colonies formed in five independent Petri dishes at 15 dpi and using ImageJ software (Schneider et al., 2012). The mycelial diameter values were divided by two to generate the radial growth (mm) values.

The degree of melanization of each tested *Z. tritici* strain was estimated from at least 40 colonies formed on PDA plates. We used a macro developed in the ImageJ (Schneider et al., 2012), which scores the mean gray value of each individual colony. Gray values range from 0 to 255, with 0 representing black and 255 representing white. The mean gray values of each strain over time were plotted in a boxplot using ggplot2 package from R (Wickham, 2009).

The impact on the cell integrity caused by the deletion of *ZtSo* was verified by exposing blastospores of *ΔZtKu70, ΔZtSo,* and *ΔZtSo*-comp to nine different stress conditions, including different temperatures (18°C and 27°C), oxidative stress (0.5, and 1 mM of hydrogen peroxide – H_2_O_2_), osmotic stress (1M sodium chloride - NaCl and 1M sorbitol), cell wall stress (2 mg/mL congo red and 10 μg/mL calcofluor white - CFW), and plasma membrane stress (0.01% sodium dodecyl sulfate – SDS). Spore suspensions of each strain were serial diluted to 4×10^4^, 4×10^5^, 4×10^6^, and 4×10^7^ blastospores/mL and drops of 3.5 μL were plated on five independent PDA plates containing the mentioned stresses and incubated at 18°C. Colony phenotypes were assessed by digital images taken at 5 dpi.

### Virulence assay and pycnidia formation *in vitro*

We used five different winter cultivars of wheat (*Triticum aestivum L.*) based on their susceptibility or resistance to *Z. tritici*, as described in the Swiss granum website (https://www.swissgranum.ch/documents/741931/1152834/LES_Winterweizen_2020.pdf/e624760c-8329-e11d-7e53-afbec9146156). Drifter, Claro and Runal are classified as susceptible cultivars. Arina is considered intermediate and Titlis and Camedo are described as resistant cultivars to *Z. tritici*. Seeding, greenhouse and plant growth conditions, inoculum preparation and plant inoculation followed the same procedures described by Meile et al. (2018). To estimate the percentage of leaf-covered by lesions (PLACL) and pycnidia formation, we harvested the second leaves of Drifter plants inoculated with *ΔZtKu70*, *ΔZtSo* or *ΔZtSo*-comp strains at 8, 10, 11, 12, 14, 16, and 21 dpi. For the rest of the cultivars, infected leaves were harvested only at 14 and 21 dpi to check for pycnidia formation. Leaves were mounted on a paper sheet, scanned with a flatbed scanner, and analyzed using automated image analysis (Stewart *et al.*, 2016). Data analysis and plotting were performed using ggplot2 package (Wickham, 2009).

The defect in pycnidia formation was confirmed by plating blastospores of *ΔZtKu70*, *ΔZtSo* and *ΔZtSo*-comp strains onto wheat extract agar medium (50 g/L blended 21 days-old wheat leaves cv. Drifter, 10 g/L agar) and incubated under UV-A light (16:8 light:dark cycle) up to 40 days at 18°C.

### Confocal laser-scanning microscopy of infected wheat leaves

To assess the impact of *ZtSo* deletion on fungal fitness during host colonization, we inoculated wheat plants with 1E4_GFP_ and other two independent GFP-tagged *ΔZtSo* mutants. Infected leaves were harvested at 6, 7, 8, 9, 10, 11, and 12 dpi and checked for developmental stages of asexual fruiting bodies. Microscopy was conducted using Zeiss LSM 780 inverted laser-scanning microscope with ZEN Black 2012 software. An argon laser at 500 nm was used to exited GFP fluorescence and chloroplast autofluorescence with an emission wavelength of 490-535 nm and 624-682 nm, respectively. Analyses, visualization, and processing of image z-stacks were performed using ImageJ software (Schneider et al., 2012).

## ACKNOWNLEDGEMENTS

This study was financed in part by the Coordenação de Aperfeiçoamento de Pessoal de Nível Superior – Brasil (CAPES) – Finance Code 001. We acknowledge Marc-Henri Lebrun from the National Institute of Agricultural Research (INRA) in France for kindly providing the MAP kinase mutants used in this study. We thank Dominik Vetsch for his help during the plant assays using different wheat cultivars.

## Supporting Information

**Table S1.** List of primers used in this study for cloning, sequencing and knock-out mutants confirmation.

**Figure S1.** Vegetative hyphal fusion occurs during epiphytic growth on wheat leaves. Co-infection of wheat plants with blastospores (A) or pycnidiospores (B) expressing either the cytoplasmic green fluorescent protein (GFP) or the red-fluorescent protein (mCherry) resulted in hyphal fusions and cytoplasm streaming of both fluorescent proteins after 48 hours of infection. Hyphal fusion during epiphytic colonization may assist the fungus to create an interconnected network supporting its establishment on the leaf surface before host penetration.

**Figure S2.** Scheme showing *ZtS*o gene and protein sequences and its phylogenetic relationship among Ascomycete species. (A) Illustration demonstrates the *ZtSo* gene locus and its protein sequencing containing the Atrophin 1, WW and PhoD as protein domains. (B) The alignment of the WW protein-protein interaction domain including the PPLP motif of 41 different fungal species. Red boxes surround the two conserved tryptophan residues spaced 22 amino acids apart. (C) Phylogenetic analysis grouped the orthologues of the *ZtSo* gene onto three groups based on fungal Classes (Dothideomycetes, Sordariomycetes, and Chaetothyriomycetes together with Eurotiomycetes), independently whether they were parasites, mutualists or saprotrophs. Three members of Basidiomycetes were used as an outgroup.

**Figure S3.** Cytoplasmic streaming between *ΔZtKu70* and the GFP-tagged 1E4 strain. Blastospores of *ΔZtKu70* and 1E4_(GFP)_ were co-inoculated on water agar (WA) plates, a hyphal fusion-inducing condition. After 40 hours of incubation, fusion bridges were observed between *ΔZtKu70* and 1E4_(GFP)_ strains. The detection of the green fluorescent protein in the cytoplasm of the recipient hypha *ΔZtKu70* confirms the cytoplasmic streaming between the two fused individuals (panel 1). Black asterisk points to the non-fluorescent *ΔZtKu70* spore before to hyphal fusion. White triangle indicates the fusion point between the *ΔZtKu70* and 1E4_(GFP)_ strains.

**Figure S4.** Cytoplasmic streaming between *ΔZtSo*-comp and GFP-tagged 1E4 strain. Blastospores of *ΔZtSo*-comp and 1E4_(GFP)_ were co-inoculated on water agar (WA) plates, a hyphal fusion-inducing condition. After 40 hours of incubation, fusion bridges were observed between *ΔZtSo*-comp and 1E4_(GFP)_ strains. The detection of the green fluorescent protein in the cytoplasm of the recipient hyphae *ΔZtSo*-comp confirm the cytoplasmic streaming between the fused individuals (panel 1). Black asterisks point to the non-fluorescent *ΔZtSo*-comp spore before to hyphal fusion. White triangles indicate the fusion points between the *ΔZtSo*-comp and 1E4_(GFP)_ strains.

**Figure S5.** Co-inoculation of *ΔZtSo* and GFP-tagged 1E4 strain confirms the failure of the *ΔZtSo* mutant to undergo hyphal fusion. Blastospores of *ΔZtSo* and 1E4_(GFP)_ were co-inoculated on water agar (WA) plates, a hyphal fusion-inducing condition. After 40 hours of incubation, fusion bridges were only observed between 1E4_(GFP)_ germinating spores (panel 1). Fluorescent green protein was never detected on the cytoplasm of *ΔZtSo* cells (panels 1, 2 and 3). The filamentous hyphae of *ΔZtSo* mutant could grow in parallel with those hyphae from the 1E4_(GFP)_, but they never undergo hyphal fusion (panels 2 and 3), demonstrating that *ZtSo* is required in both fusion partners for the establishment of the fungal communication required for perception and attraction before fusion. Black asterisks point to the non-fluorescent *ΔZtSo* spores.

**Figure S6.** MAP kinase-encoding *ZtSlt2* and *ZtFus3* genes are required for anastomosis in *Zymoseptoria tritici*. Hyphal fusions were regularly found for the wild-type strain (IPO323). However, deletion of *ZtSlt2* (orthologous to MAK-1) or *ZtFus3* (orthologous to MAK-2) resulted in fusion-defective mutants, probably due to the disruption of the oscillatory recruitment of both MAP kinase modules required for cell-to-cell communication and fusion, as described for *Neurospora crassa* (Fleissner et al., 2009). The defective phenotype was restored in the complemented *ΔZtSlt2*-comp strain. White triangles point to self-fusion events.

**Figure S7.** Effect of *ZtSo* deletion on the vegetative fungal growth. (A) The *ΔZtSo* mutant exhibited a similar colony morphology than *ΔZtKu70* and *ΔZtSo*-comp strains. (B) Light microscopy of colony edges showed a dense hyphal-thickened margin for *ΔZtKu70* and *ΔZtSo*-comp colonies, whereas the *ΔZtSo* mutant exhibited only a few filamentous at the colony periphery. Dashed squares point to the localization of microscope images. (C) Thought none morphological differences were observed for the tested *Zymoseptoria tritici* strains, the fusion defective *ΔZtSo* mutant had a slight, but significant reduction of its radial growth (mm) compared to *ΔZtKu70* and *ΔZtSo*-comp, when grown on a nutrient-limited medium. At least 40 colonies of each tested strains were evaluated. Two and three stars indicate a p-value <0.005 and <0.0005, respectively. (D) None morphological differences were noticed for the blastospores produced by *ΔZtKu70, ΔZtSo* or *ΔZtSo*-comp strains. (E) *ΔZtSo* mutant grew faster and had greater radial growth overtime than the *ΔZtKu70* and *ΔZtSo*-comp strains, when incubated on a nutrient-rich medium. Bars represent standard errors of the radial growth (mm) of at least 40 colonies. Different letters on the top of the bars indicate a significant difference among the tested strains according to the Analysis of Variance (ANOVA). The notch displays a 95% confidence interval of the median. Open circles represent the outlier values of each strain.

**Figure S8.** Percentage of leaves cover by lesions (PLACL) produced by *ΔZtKu70, ΔZtSo,* and *ΔZtSo*-comp strains. The second leaves of the wheat plants cv. Drifter infected with the *Zymoseptoria tritici* strains were harvested at different days post-inoculation (dpi). Bars represent standard errors of PLACL values of at least six infected leaves. Different letters on the top of the bars indicate significant difference among the tested strains according to the Analysis of Variance (ANOVA). Black circles represent the outlier data points.

**Figure S9.** *ΔZtSo* and *ΔZtKu70* strains does not vary in pathogenicity, except for the failure of *ΔZtSo* mutant to undergo asexual reproduction. Five different winter cultivars were infected with *ΔZtKu70* or *ΔZtSo* strains and evaluated at 14- and 21 days post-inoculation (dpi) for host damage and pathogen reproduction, respectively. Runal and Claro are classified as susceptible cultivars. Arina is considered intermediate, whereas Titlis and Camedo are described as resistant cultivars to *Z. tritici.* On the left panel, *ΔZtKu70* and *ΔZtSo* showed a comparable host damage for the Runal, Claro and Titlis cultivars at 14 dpi. Both strains were avirulent in Arina and Camedo. On the right panel, the asexual reproductive structures were observed within the necrosis of those plants inoculated with the *ΔZtKu70* strain at 21 dpi. None pycnidium was observed for plants inoculated with the *ΔZtSo* mutant, demonstrating that the failure to undergo asexual reproduction is associated with the disruption of *ZtSo per se* than a cultivar-specific interaction.

**Figure S10.** Schematic demonstration of hyphal penetration, substomatal colonization and initial stages of pycnidial development. Susceptible wheat cv. Drifter was inoculated with the fluorescent *Zymoseptoria tritici* 1E4_GFP_ (wild-type) and *ΔZtSo*_GFP_ strains and monitored by confocal microscopy at different days post-infection (dpi). For later time points (9 and 12 dpi), please see Fig.7. At 6 dpi, the epiphytic filamentous hyphae penetrate the stomata and enter first in the substomatal cavity. At 7 dpi, initiate the intracellular hyphal colonization, which the filamentous surrounding the stomatal guard cells produce specialized knots from where secondary hyphae emerge and germinate. Up to this point, none morphological difference for hyphal extension and intracellular hyphal colonization was noticed between 1E4_GFP_ and *ΔZtSo*_GFP_ strains. At 8 dpi, the secondary hyphae from 1E4_GFP_ strain fuse with another nearby hypha (represented by black circles), creating an interconnected network in the sub-stomatal cavity. Unlike, the secondary hyphae from *ΔZtSo*_GFP_ kept extending as individual filamentous and none indication of anastomosis was observed until this developmental stage.

**Figure S11.** Description of functional characterizations performed in this study. (A) *ZtKu70* gene was disrupted by the geneticin resistance cassette (*gen*) via homologous recombination in the 1E4 wild-type (WT) strains and generating the 1E4*ΔZtKu70* mutant. Next, the *ZtSo* gene was knocked-out by the hygromycin resistance cassette (*hph*) via homologous recombination in the genetic background of the 1E4*ΔZtKu70* mutant, generating the double mutant 1E4*ΔZtKu70ΔZtSo*. Later, the *ZtSo* gene was re-introduced *in locus* in the previous double mutant 1E4*ΔZtKu70ΔZtSo* using the selective nourseothricin resistance gene (*nat*) and generating the 1E4*ΔZtKu70ΔZtSo*-comp mutant. (B) Agarose gel shows the PCR fragments at the expected sizes of 3’954 base pairs (bp) or 1’626 bp, confirming the presence of the *ZtSo* native gene or the hygromycin resistance gene, respectively. (C) Agarose gel shows a PCR fragment of 286 bp confirming the presence of the nourseothricin resistance gene only in the 1E4*ΔZtKu70ΔZtSo*-comp mutant.

## REFERENCES

Aldabbous, M. S., Roca, M. G., Stout, A., Huang, I. C., Read, N. D. and Free, S. J. (2010) The ham-5, rcm-1 and rco-1 genes regulate hyphal fusion in Neurospora crassa. Microbiology, 156, 2621–2629.

Biella, S., Smith, M. L., Aist, J. R., Cortesi, P. and Milgroom, M. G. (2002) Programmed cell death correlates with virus transmission in a filamentous fungus. Proc Biol Sci, 269, 2269–2276.

Bloemendal, S. and Kuck, U. (2013) Cell-to-cell communication in plants, animals, and fungi: a comparative review. Naturwissenschaften, 100, 3–19.

Bork, P. and Sudol, M. (1994) The WW domain - a signalling site in dystrophin? Trends in Biochemical Science, 19, 531–533.

Castanera, R., Lopez-Varas, L., Borgognone, A., LaButti, K., Lapidus, A., Schmutz, J., et al. (2016) Transposable Elements versus the Fungal Genome: Impact on Whole-Genome Architecture and Transcriptional Profiles. PLoS Genet, 12, e1006108.

Chagnon, P. L. (2014) Ecological and evolutionary implications of hyphal anastomosis in arbuscular mycorrhizal fungi. FEMS Microbiol Ecol, 88, 437–444.

Charlton, N. D., Shoji, J. Y., Ghimire, S. R., Nakashima, J. and Craven, K. D. (2012) Deletion of the fungal gene soft disrupts mutualistic symbiosis between the grass endophyte Epichloe festucae and the host plant. Eukaryot Cell, 11, 1463–1471.

Chu, Y. M., Jeon, J. J., Yea, S. J., Kim, Y. H., Yun, S. H., Lee, Y. W., et al. (2002) Double-stranded RNA mycovirus from Fusarium graminearum. Appl Environ Microbiol, 68, 2529–2534.

Cottier, F. and Muhlschlegel, F. A. (2012) Communication in fungi. Int J Microbiol, 2012, 351832.

Cousin, A., Mehrabi, R., Guilleroux, M., Dufresne, M., T, V. D. L., Waalwijk, C., et al. (2006) The MAP kinase-encoding gene MgFus3 of the non-appressorium phytopathogen Mycosphaerella graminicola is required for penetration and in vitro pycnidia formation. Mol Plant Pathol, 7, 269–278.

Craven, K. D., Velez, H., Cho, Y., Lawrence, C. B. and Mitchell, T. K. (2008) Anastomosis is required for virulence of the fungal necrotroph Alternaria brassicicola. Eukaryot Cell, 7, 675–683.

Dean, R., Van Kan, J. A., Pretorius, Z. A., Hammond-Kosack, K. E., Di Pietro, A., Spanu, P. D., et al. (2012) The Top 10 fungal pathogens in molecular plant pathology. Mol Plant Pathol, 13, 414–430.

Deng, Y. Z., Zhang, B., Chang, C., Wang, Y., Lu, S., Sun, S., et al. (2018) The MAP Kinase SsKpp2 Is Required for Mating/Filamentation in Sporisorium scitamineum. Front Microbiol, 9, 2555.

Dhillon, B., Gill, N., Hamelin, R. C. and Goodwin, S. B. (2014) The landscape of transposable elements in the finished genome of the fungal wheat pathogen Mycosphaerella graminicola. BMC Genomics, 15, 1132.

Endler, J. A. (1993) Some general comments on the evolution and design of animal communication systems. Philosophical Transactions of the Royal Society B, 340, 215–225.

Engh, I., Wurtz, C., Witzel-Schlomp, K., Zhang, H. Y., Hoff, B., Nowrousian, M., et al. (2007) The WW domain protein PRO40 is required for fungal fertility and associates with Woronin bodies. Eukaryot Cell, 6, 831–843.

Fischer, M. S. and Glass, N. L. (2019) Communicate and Fuse: How Filamentous Fungi Establish and Maintain an Interconnected Mycelial Network. Front Microbiol, 10, 619.

Fleissner, A. and Herzog, S. (2016) Signal exchange and integration during self-fusion in filamentous fungi. Semin Cell Dev Biol, 57, 76–83.

Fleissner, A., Leeder, A. C., Roca, M. G., Read, N. D. and Glass, N. L. (2009) Oscillatory recruitment of signaling proteins to cell tips promotes coordinated behavior during cell fusion. Proc Natl Acad Sci U S A, 106, 19387–19392.

Fleissner, A., Sarkar, S., Jacobson, D. J., Roca, M. G., Read, N. D. and Glass, N. L. (2005) The so locus is required for vegetative cell fusion and postfertilization events in Neurospora crassa. Eukaryot Cell, 4, 920–930.

Fones, H. and Gurr, S. (2015) The impact of Septoria tritici Blotch disease on wheat: An EU perspective. Fungal Genet Biol, 79, 3–7.

Francisco, C. S., Ma, X., Zwyssig, M. M., McDonald, B. A. and Palma-Guerrero, J. (2019) Morphological changes in response to environmental stresses in the fungal plant pathogen Zymoseptoria tritici. Sci Rep, 9, 9642.

Friesen, T. L., Stukenbrock, E. H., Liu, Z., Meinhardt, S., Ling, H., Faris, J. D., et al. (2006) Emergence of a new disease as a result of interspecific virulence gene transfer. Nat Genet, 38, 953–956.

Fu, C., Ao, J., Dettmann, A., Seiler, S. and Free, S. J. (2014) Characterization of the Neurospora crassa cell fusion proteins, HAM-6, HAM-7, HAM-8, HAM-9, HAM-10, AMPH-1 and WHI-2. PLoS One, 9, e107773.

Fu, C., Iyer, P., Herkal, A., Abdullah, J., Stout, A. and Free, S. J. (2011) Identification and characterization of genes required for cell-to-cell fusion in Neurospora crassa. Eukaryot Cell, 10, 1100–1109.

Gillam, E. (2011) An introduction to animal communication. Nature Education Knowledge 3, 70.

Glass, N. L., Jacobson, D. J. and Shiu, P. K. T. (2000) The genetics of hyphal fusion and vegetative incompatibility in filamentous ascomycete fungi. Annual Review of Genetics, 34, 165–186.

Goddard, M. R. and Burt, A. (1999) Recurrent invasion and extinction of a selfish gene. PNAS, 96, 13880–13885.

Gohari, A. M., Mehrabi, R., Robert, O., Ince, I. A., Boeren, S., Schuster, M., et al. (2014) Molecular characterization and functional analyses of ZtWor1, a transcriptional regulator of the fungal wheat pathogen Zymoseptoria tritici. Mol Plant Pathol, 15, 394–405.

Hagiwara, D., Sakamoto, K., Abe, K. and Gomi, K. (2016) Signaling pathways for stress responses and adaptation in Aspergillus species: stress biology in the post-genomic era. Biosci Biotechnol Biochem, 80, 1667–1680.

He, C., Rusu, A. G., Poplawski, A. M., Irwin, J. A. G. and Manners, J. M. (1998) Transfer of a Supernumerary Chromosome Between Vegetatively Incompatible Biotypes of the Fungus Colletotrichum gloeosporioides. Genetics Society of America, 150, 1459–1466.

Hickey, P. C., Jacobson, D. J., Read, N. D. and Glass, N. L. (2002) Live-cell imaging of vegetative hyphal fusion in Neurospora crassa. Fungal Genetics and Biology, 37, 109–119.

Ihrmark, K., Johannesson, H., Stenström, E. and Stenlid, J. (2002) Transmission of double-stranded RNA in Heterobasidion annosum. Fungal Genetics and Biology, 36, 147–154.

Ishikawa, F. H., Souza, E. A., Read, N. D. and Roca, M. G. (2010) Live-cell imaging of conidial fusion in the bean pathogen, Colletotrichum lindemuthianum. Fungal Biol, 114, 2–9.

Kema, G. H. and van Silfhout, C. H. (1997) Genetic Variation for Virulence and Resistance in the Wheat-Mycosphaerella graminicola Pathosystem III. Comparative Seedling and Adult Plant Experiments. Phytopathology, 87, 266–272.

Kema, G. H. J., van der Lee, T. A. J., Mendes, O., Verstappen, E. C. P., Lankhorst, R. K., Sandbrink, H., et al. (2008) Large-Scale Gene Discovery in the Septoria Tritici Blotch Fungus Mycosphaerella graminicola with a Focus on In Planta Expression. Molecular Plant-Microbe Interactions, 21, 1249–1260.

Khang, C. H., Berruyer, R., Giraldo, M. C., Kankanala, P., Park, S. Y., Czymmek, K., et al. (2010) Translocation of Magnaporthe oryzae effectors into rice cells and their subsequent cell-to-cell movement. Plant Cell, 22, 1388–1403.

Kilaru, S., Schuster, M., Ma, W. and Steinberg, G. (2017) Fluorescent markers of various organelles in the wheat pathogen Zymoseptoria tritici. Fungal Genet Biol, 105, 16–27.

Krishnan, P., Meile, L., Plissonneau, C., Ma, X., Hartmann, F. E., Croll, D., et al. (2018) Transposable element insertions shape gene regulation and melanin production in a fungal pathogen of wheat. BMC Biol, 16, 78.

Larsson, A. (2014) AliView: a fast and lightweight alignment viewer and editor for large datasets. Bioinformatics, 30, 3276–3278.

Lendenmann, M. H., Croll, D., Stewart, E. L. and McDonald, B. A. (2014) Quantitative trait locus mapping of melanization in the plant pathogenic fungus Zymoseptoria tritici. G3 (Bethesda), 4, 2519–2533.

Leng, Y. and Zhong, S. (2015) The Role of Mitogen-Activated Protein (MAP) Kinase Signaling Components in the Fungal Development, Stress Response and Virulence of the Fungal Cereal Pathogen Bipolaris sorokiniana. PLoS One, 10, e0128291.

Levin, D. E. (2005) Cell wall integrity signaling in Saccharomyces cerevisiae. Microbiol Mol Biol Rev, 69, 262–291.

Liu, W., Soulie, M. C., Perrino, C. and Fillinger, S. (2011) The osmosensing signal transduction pathway from Botrytis cinerea regulates cell wall integrity and MAP kinase pathways control melanin biosynthesis with influence of light. Fungal Genet Biol, 48, 377–387.

Maddi, A., Dettman, A., Fu, C., Seiler, S. and Free, S. J. (2012) WSC-1 and HAM-7 are MAK-1 MAP kinase pathway sensors required for cell wall integrity and hyphal fusion in Neurospora crassa. PLoS One, 7, e42374.

Maruyama, J., Escano, C. S. and Kitamoto, K. (2010) AoSO protein accumulates at the septal pore in response to various stresses in the filamentous fungus Aspergillus oryzae. Biochem Biophys Res Commun, 391, 868–873.

Mat Razali, N., Cheah, B. H. and Nadarajah, K. (2019) Transposable Elements Adaptive Role in Genome Plasticity, Pathogenicity and Evolution in Fungal Phytopathogens. Int J Mol Sci, 20.

McDonald, M. C., Taranto, A. P., Hill, E., Schwessinger, B., Liu, Z., Simpfendorfer, S., et al. (2019) Transposon-Mediated Horizontal Transfer of the Host-Specific Virulence Protein ToxA between Three Fungal Wheat Pathogens. MBio, 10.

Mehrabi, R., Bahkali, A. H., Abd-Elsalam, K. A., Moslem, M., Ben M’barek, S., Gohari, A. M., et al. (2011) Horizontal gene and chromosome transfer in plant pathogenic fungi affecting host range. FEMS Microbiol Rev, 35, 542–554.

Mehrabi, R., Ben M’Barek, S., van der Lee, T. A., Waalwijk, C., de Wit, P. J. and Kema, G. H. (2009) G(alpha) and Gbeta proteins regulate the cyclic AMP pathway that is required for development and pathogenicity of the phytopathogen Mycosphaerella graminicola. Eukaryot Cell, 8, 1001–1013.

Mehrabi, R., van der Lee, T., Waalwijk, C. and Kema, G. H. (2006) MgSlt2, a cellular integrity MAP kinase gene of the fungal wheat pathogen Mycosphaerella graminicola, is dispensable for penetration but essential for invasive growth. Molecular Plant-Microbe Interactions, 19, 389–398.

Meile, L., Croll, D., Brunner, P. C., Plissonneau, C., Hartmann, F. E., McDonald, B. A., et al. (2018) A fungal avirulence factor encoded in a highly plastic genomic region triggers partial resistance to septoria tritici blotch. New Phytol.

Mendiburu, F. D. (2015) Agricolae: Statistical Procedures for Agricultural Research. R Package Version 1.2-3.

Nuss, D. L. (2005) Hypovirulence: mycoviruses at the fungal-plant interface. Nat Rev Microbiol, 3, 632–642.

Ord, T. J. and Garcia-Porta, J. (2012) Is sociality required for the evolution of communicative complexity? Evidence weighed against alternative hypotheses in diverse taxonomic groups. Philos Trans R Soc Lond B Biol Sci, 367, 1811–1828.

Pandey, A., Roca, M. G., Read, N. D. and Glass, N. L. (2004) Role of a mitogen-activated protein kinase pathway during conidial germination and hyphal fusion in Neurospora crassa. Eukaryot Cell, 3, 348–358.

Park, G., Pan, S. and Borkovich, K. A. (2008) Mitogen-activated protein kinase cascade required for regulation of development and secondary metabolism in Neurospora crassa. Eukaryot Cell, 7, 2113–2122.

Pearson, M. N., Beever, R. E., Boine, B. and Arthur, K. (2009) Mycoviruses of filamentous fungi and their relevance to plant pathology. Mol Plant Pathol, 10, 115–128.

Plonka, P. M. and Grabacka, M. (2006) Melanin synthesis in microorganisms — biotechnological and medical aspects. Acta Biochimica Polonica, 53, 429–443.

Prados Rosales, R. C. and Di Pietro, A. (2008) Vegetative Hyphal Fusion Is Not Essential for Plant Infection by Fusarium oxysporum. Eukaryotic Cell, 7, 162.

Read, N. D., Fleissner, A., Roca, M. G. and Glass, N. L. (2010) Hyphal Fusion. In: Cellular abd Molecular Biology of Filamentous Fungi. (Borkovich, K. A. and Edolle, D., eds.). Washington DC: American Society of Microbiology, pp. 260–273.

Read, N. D., Goryachev, A. B. and Lichius, A. (2012) The mechanistic basis of self-fusion between conidial anastomosis tubes during fungal colony initiation. Fungal Biology Reviews, 26, 1–11.

Read, N. D., Lichius, A., Shoji, J. Y. and Goryachev, A. B. (2009) Self-signalling and self-fusion in filamentous fungi. Curr Opin Microbiol, 12, 608–615.

Roca, G. M., Read, N. D. and Wheals, A. E. (2005a) Conidial anastomosis tubes in filamentous fungi. FEMS Microbiol Lett, 249, 191–198.

Roca, M. G., Arlt, J., Jeffree, C. E. and Read, N. D. (2005b) Cell biology of conidial anastomosis tubes in Neurospora crassa. Eukaryot Cell, 4, 911–919.

Roca, M. G., Davide, L. C., Davide, L. M., Mendes-Costa, M. C., Schwan, R. F. and Wheals, A. E. (2004) Conidial anastomosis fusion between Colletotrichum species. Mycol Res, 108, 1320–1326.

Rodrigues, M. L., Nakayasu, E. S., Oliveira, D. L., Nimrichter, L., Nosanchuk, J. D., Almeida, I. C., et al. (2008) Extracellular vesicles produced by Cryptococcus neoformans contain protein components associated with virulence. Eukaryot Cell, 7, 58–67.

Roper, M., Ellison, C., Taylor, J. W. and Glass, N. L. (2011) Nuclear and genome dynamics in multinucleate ascomycete fungi. Curr Biol, 21, R786–793.

Saupe, S. J. (2000) Molecular Genetics of Heterokaryon Incompatibility in Filamentous Ascomycetes. Microbiology and Molecular Biology Reviews, 64, 489–502.

Schneider, C. A., Rasband, W. S. and Eliceiri, K. W. (2012) NIH Image to ImageJ: 25 years of image analysis. Nature Methods, 9, 671.

Shrout, J. D., Tolker-Nielsen, T., Givskov, M. and Parsek, M. R. (2011) The contribution of cell-cell signaling and motility to bacterial biofilm formation. MRS Bull, 36, 367–373.

Sidhu, Y. S., Cairns, T. C., Chaudhari, Y. K., Usher, J., Talbot, N. J., Studholme, D. J., et al. (2015) Exploitation of sulfonylurea resistance marker and non-homologous end joining mutants for functional analysis in Zymoseptoria tritici. Fungal Genet Biol, 79, 102–109.

Silva, B. M., Prados-Rosales, R., Espadas-Moreno, J., Wolf, J. M., Luque-Garcia, J. L., Goncalves, T., et al. (2014) Characterization of Alternaria infectoria extracellular vesicles. Med Mycol, 52, 202–210.

Simonin, A., Palma-Guerrero, J., Fricker, M. and Glass, N. L. (2012) Physiological significance of network organization in fungi. Eukaryot Cell, 11, 1345–1352.

Stewart, E. L., Hagerty, C. H., Mikaberidze, A., Mundt, C. C., Zhong, Z. and McDonald, B. A. (2016) An Improved Method for Measuring Quantitative Resistance to the Wheat Pathogen Zymoseptoria tritici Using High-Throughput Automated Image Analysis. Phytopathology, 106, 782–788.

Tamura, K., Stecher, G., Peterson, D., Filipski, A. and Kumar, S. (2013) MEGA6: Molecular Evolutionary Genetics Analysis version 6.0. Mol Biol Evol, 30, 2725–2729.

Teichert, I., Steffens, E. K., Schnass, N., Franzel, B., Krisp, C., Wolters, D. A., et al. (2014) PRO40 is a scaffold protein of the cell wall integrity pathway, linking the MAP kinase module to the upstream activator protein kinase C. PLoS Genet, 10, e1004582.

Temporini, E. D. and VanEtten, H. D. (2004) An analysis of the phylogenetic distribution of the pea pathogenicity genes of Nectria haematococca MPVI supports the hypothesis of their origin by horizontal transfer and uncovers a potentially new pathogen of garden pea: Neocosmospora boniensis. Curr Genet, 46, 29–36.

Valiante, V. (2017) The Cell Wall Integrity Signaling Pathway and Its Involvement in Secondary Metabolite Production. J Fungi (Basel), 3.

Valiante, V., Macheleidt, J., Foge, M. and Brakhage, A. A. (2015) The Aspergillus fumigatus cell wall integrity signaling pathway: drug target, compensatory pathways, and virulence. Front Microbiol, 6, 325.

van Gestel, J., Nowak, M. A. and Tarnita, C. E. (2012) The evolution of cell-to-cell communication in a sporulating bacterium. PLoS Comput Biol, 8, e1002818.

Wickham, H. (2009) ggplot2: Elegant Graphics for Data Analysis. New York: Use R. Springer-Verlag.

Wilson, E. O. (1975) Sociobiology: the new synthesis. Harvard University Press.

Wongsuk, T., Pumeesat, P. and Luplertlop, N. (2016) Fungal quorum sensing molecules: Role in fungal morphogenesis and pathogenicity. J Basic Microbiol, 56, 440–447.

Xiang, Q., Rasmussen, C. and Glass, N. L. (2002) The ham-2 Locus, Encoding a Putative Transmembrane Protein, Is Required for Hyphal Fusion in Neurospora crassa. Genetics, 160, 169–180.

Yago, J. I., Lin, C. H. and Chung, K. R. (2011) The SLT2 mitogen-activated protein kinase-mediated signalling pathway governs conidiation, morphogenesis, fungal virulence and production of toxin and melanin in the tangerine pathotype of Alternaria alternata. Mol Plant Pathol, 12, 653–665.

Zelikovitch, N., Eyal, Z., Ben-Zyi, B. and Koltin, Y. (1990) Double-stranded RNA mycoviruses in Septoria tritici. Mycol Res, 94, 590–594.

Zhan, J., Kema, G. H., Waalwijk, C. and McDonald, B. A. (2002) Distribution of mating type alleles in the wheat pathogen Mycosphaerella graminicola over spatial scales from lesions to continents. Fungal Genetics and Biology, 36, 128–136.

Zhao, X., Mehrabi, R. and Xu, J. R. (2007) Mitogen-activated protein kinase pathways and fungal pathogenesis. Eukaryot Cell, 6, 1701–1714.

